# Cardiomyocyte vulnerability to lamin polymer disruption revealed by saturation mutagenesis

**DOI:** 10.64898/2026.09.17.752442

**Authors:** Jessica Mella, Abigail Hein, Navraj Lally, Julia Conrad, Cathy Yang, Andrew P. Landstrom, Vasanth Vedantham, Willow Coyote-Maestas, Abigail Buchwalter

**Affiliations:** Cardiovascular Research Institute, University of California, San Francisco, USA; Tetrad Graduate Program, University of California, San Francisco, USA; Biomedical Sciences Graduate Program, University of California, San Francisco, USA; Department of Biochemistry, University of California, San Francisco, USA; Department of Pediatrics, Division of Cardiology, Duke University School of Medicine, Durham, North Carolina, USA; Department of Pediatrics, Division of Cardiology, Children’s Hospital of Philadelphia and Perelman School of Medicine, University of Pennsylvania, Philadelphia, Pennsylvania, USA; Cardiology Division, Department of Medicine, University of California, San Francisco, USA; Department of Bioengineering and Therapeutic Sciences, University of California, San Francisco, USA; Chan Zuckerberg Biohub, San Francisco, USA; Quantitative Biosciences Institute, University of California, San Francisco, USA

## Abstract

Hundreds of mutations to the broadly expressed *LMNA* gene cause disease primarily within cardiac, muscular, and adipose tissues^1^. Tissue-specific pathogenesis arises when mutant protein dysfunction collides with the unique demands of a specific cell type. Here, we decipher the cell-type-specific consequences of ∼15,000 *LMNA* mutations by completing the first saturation mutagenesis screens in human induced pluripotent cells (hiPSCs) and hiPSC-derived cardiomyocytes using our newly developed <u>s</u>ingle <u>l</u>arge serine <u>i</u>ntegrase <u>c</u>assette <u>e</u>xchange (SLICE) platform. We find that destabilization is a predominant consequence of pathogenic *LMNA* mutations, is selected against in human populations, and is associated with cardiomyopathy. Mutation sensitivity maps reveal Lamin A quality control at both the subunit and multimer level, resolve lateral and head-to-tail polymerization interfaces, and uncover a convergence between disruption of lamin polymer assembly and pathogenesis. Uniquely in cardiomyocytes, lamin A polymer assembly defects drive profound protein loss, nuclear abnormalities, and cellular toxicity, explaining the origins of cardiac specificity in laminopathy syndromes.

## Main Text

In human disease, the variable effects of missense mutations in protein-coding sequences shape both disease mechanisms and responses to therapy. Loss-of-function (LOF) mutations frequently disrupt protein folding and prevent normal function, while gain-of-function (GOF) mutations often have milder effects on protein structure but can dominantly disrupt function^2^. Rational design of therapies depends on a molecular understanding of how each pathogenic mutation affects protein structure and function.

The nuclear lamina is an essential structure that protects and organizes the genome. Mutations to its constituent proteins cause “laminopathy” syndromes, the most common of which is a dilated cardiomyopathy (DCM) linked to the *LMNA* gene^1^. *LMNA*-linked DCM occurs at an estimated population frequency of ∼1 in 10,000 with a high incidence of life-threatening arrhythmias and a particularly severe disease course^3^. NCBI’s ClinVar database currently identifies more than 600 missense, nonsense, indel, and frameshift *LMNA* mutations as pathogenic or likely pathogenic, and lists nearly 900 variants of uncertain significance (VUS). For some *LMNA* mutations, the mechanism of action is clear: early nonsense truncating mutations prevent protein production from the mutant allele by destabilizing the encoding transcript and/or protein, causing LOF^4^. However, *LMNA* nonsense mutations are far outnumbered by missense mutations, and it is unclear how the vast majority of these drive disease.

The lamins are an ancient subfamily of polymer-forming intermediate filament (IF) proteins. Like all IFs, lamins are coiled-coil homodimers that self-associate into higher-order multimers. Mutation-induced misfolding could thus disrupt the unit dimer, higher-order multimers, and/or the function of the assembled lamina meshwork. As lamin polymers have not yet been structurally resolved, we have limited insight into the effects of laminopathy-linked mutations on lamin assembly and function.

While the *LMNA* gene is broadly expressed, *LMNA-*linked diseases manifest primarily within the cardiovascular system, skeletal muscle, or adipose tissue^1^. Tissue-specific pathogenicity can arise when broadly expressed mutant proteins intersect with the unique tissue physiology^5^. This unique physiology includes variation in protein quality control (pQC) networks, which are responsible for the fate of mutant proteins. The pQC machinery may detect and degrade mutant proteins or become overwhelmed, resulting in irreversible protein aggregation. Differences in the response to protein misfolding across cell types can thus shape mutation effects even for universally expressed mutant proteins^6^.

The lamina is an extremely stable and long-lived structure in post-mitotic tissues^7^. Long-lived proteins exist at an extreme proteostatic setpoint of very low synthesis and degradation rates^8^. The pQC machinery of proliferative cells may have greater capacity to degrade these proteins when they are disassembled during mitosis, while the pQC machinery of postmitotic cells may have limited opportunities to access the lamins when they are locked away within the lamina. These extreme features of lamin proteostasis may make Lamin A variably sensitive to destabilization by mutations across cell states^8^ and may contribute to the tissue-specific effects of pathogenic mutations. We recently reported that a *LMNA* mutation that drives Hutchinson-Gilford progeria syndrome (HGPS) further extends Lamin A/C protein lifetime and drives protein accumulation within the cardiovascular system^7^. It is unclear whether cardiac-specific proteostasis disruptions generalize across *LMNA* mutations, as addressing this question requires a systematic analysis of *LMNA* mutation effects in cardiac cell types.

<u>D</u>eep <u>m</u>utational <u>s</u>canning (DMS) is a potent high-throughput approach for defining how sequence variation influences protein structure, assembly, and function^9^. It is now possible to generate and phenotype comprehensive variant libraries for a protein of interest, enabling the study of thousands of mutations simultaneously. As a gene with a high mutational burden and limited genotype:phenotype relationships, *LMNA* is an ideal candidate for DMS^10^. However, to date, most DMS studies have been performed in transformed cell lines that poorly model human disease. Targeted variant assessments have been performed in hiPSC-derived cardiomyocytes^11,12^, but unbiased saturation mutagenesis in these contexts has not yet been achieved. Here, we adapt DMS to human induced pluripotent stem cells (hiPSCs) and their differentiated progeny and perform unbiased saturation mutagenesis in disease-relevant cell types for the first time. Our *LMNA* screens in hiPSCs, epicardial cells, and ventricular cardiomyocytes delineate the molecular basis of Lamin A polymerization and uncover cardiomyocyte-specific vulnerability to mutations at Lamin A polymerization interfaces. These results provide new insight into the yet unresolved lamin filament structure and the tissue specificity of *LMNA* mutation effects.

### Delivering mutation libraries with single large serine integrase cassette exchange (SLICE)

To dissect the cell type-specific effects of *LMNA* sequence variation at scale, we sought to deliver a comprehensive mutation library into hiPSCs, then differentiate these cells into cardiac lineages (Fig. 1A). DMS requires expression of a single variant per cell, which becomes more difficult with increasing library size^13^. For this reason, DMS studies are often performed in immortalized cell lines with limited relevance to human disease. Directed differentiation of hiPSCs makes disease-relevant cell types accessible, but random integration-based library delivery methods often yield variable expression levels and stochastic silencing in hiPSCs^14^. No platform that supported efficient delivery and stable, single-copy expression of DMS libraries into hiPSCs previously existed. We built SLICE (<u>s</u>ingle <u>l</u>arge serine <u>i</u>ntegrase <u>c</u>assette <u>e</u>xchange), a landing pad system for stable transgene expression. SLICE incorporates several key advantages: (1) use of the recently discovered Pa01 integrase to mediate directional insertion^15^; (2) anti-silencing elements for stable expression through differentiation^14^; and (3) positive and negative selection strategies for purifying successful integrants (Fig. 1 supplement 1A-D)^15^.

**Figure 1.**
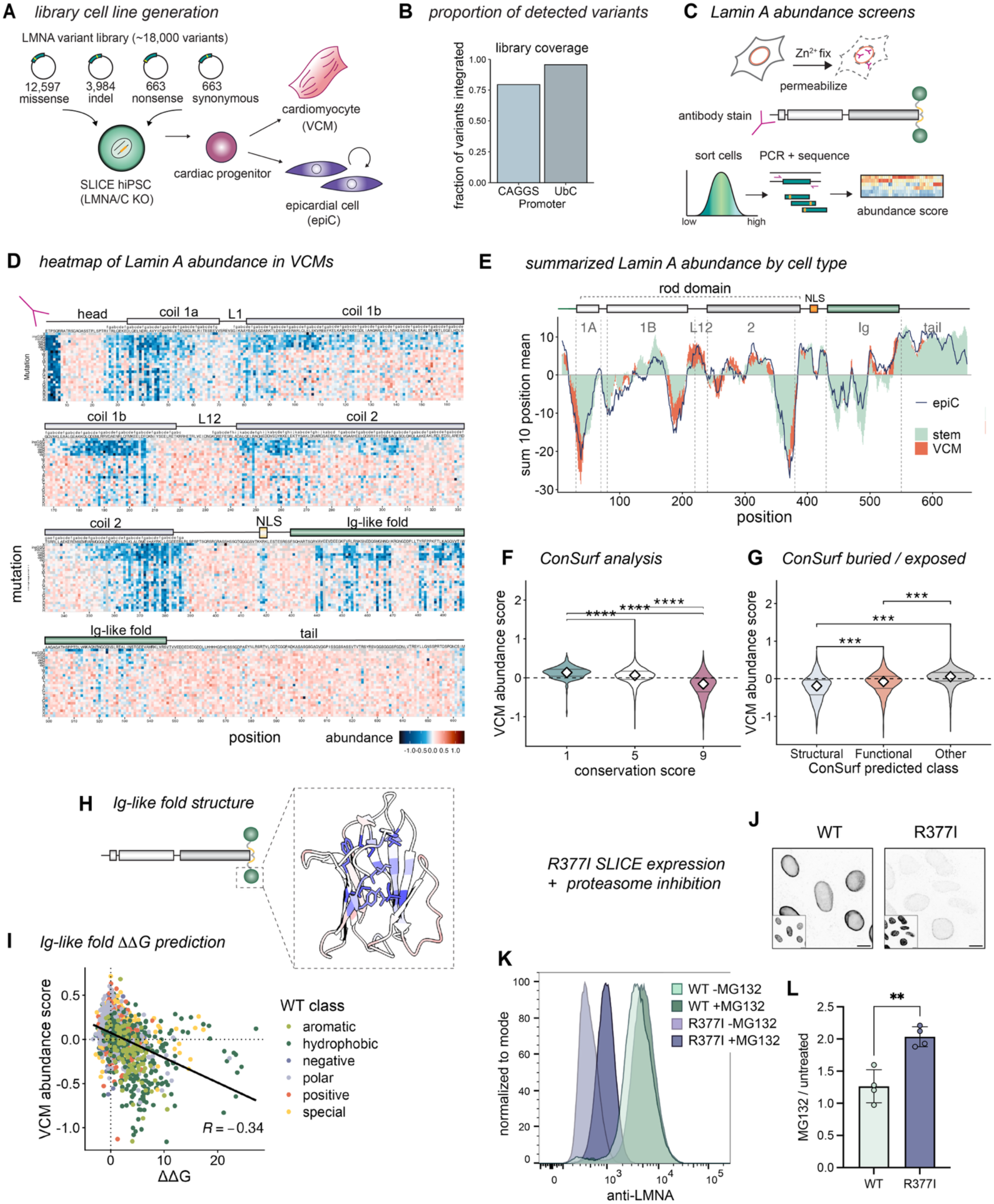
Immunolabeling screens reveal unstable lamin A variants in differentiated cardiac cells. A) Diagram of *LMNA* DMS variant library cell line generation. Pooled plasmid variant library is integrated into *LMNA* KO hiPSCs via SLICE, then differentiated into either epicardial cells (epiCs) or ventricular cardiomyocytes (VCMs). B) Percent of cloned library variants detected in hiPSCs by sequencing after SLICE integration. CAGGS promoter, library version 1 (hiPSCs, epiCs); UbC promoter, library version 2 (epiCs, VCMs). C) Diagram of immunostaining screen workflow. Cells are fixed with zinc, permeabilized, and stained with anti-LMNA/C antibodies before sorting by fluorescence intensity into 4 bins. Genomic DNA is extracted from the cells in each bin, PCR amplified from the SLICE landing pad, and sequenced to obtain variant counts. D) Heatmap of lamin A mutation abundance scores in VCMs. Full protein sequence broken into 4 stacked segments. Lamin A residue numbers along the x-axis with WT residue, heptad phase, and domain diagram at the top of the heatmap. Mutations are along the y-axis. Heatmap is colored by protein abundance, where blue indicates low abundance. N-terminal antibody epitope indicated. E) Rolling-window abundance signal along lamin A by cell type. Each trace is the rolling sum of z-scored position-mean Lilace abundance scores over 10-residue windows (1-position step), plotted at the window center, for each cell type (colors). Top, lamin A domain diagram annotating the coiled-coil segments (1a, 1b, 2), linker L12, nuclear localization signal (NLS) Ig-like fold, and tail. F) VCM abundance by conservation. Mean values at each position were plotted by ConSurf-assigned categorical score: 1 = variable, 5 = average, and 9 = conserved. \*\*\*\**P* < 0.0001 by pairwise Wilcoxon test with FDR correction. G) VCM abundance by predicted functional class. Highly conserved (ConSurf 8–9) variants predicted to be on the surface were assigned “Functional” class, while highly conserved variants predicted to be buried were assigned “Structural” class. Variants with lower conservation classified as “Other.” H) Lamin A globular Ig-like fold structure (PDB 1IFR) colored by positional mean VCM abundance score, where blue indicates low abundance. Hydrophobic, buried residues indicated by stick display. I) VCM abundance scores for each variant in the Ig-like fold plotted against FoldX-predicted ΔΔG. Color indicates chemical character of WT residues. J) Confocal images of *LMNA* KO epicardial cells expressing lamin A WT or R377I mutant from the SLICE landing pad immunostained for LMNA/C. Inset, Hoechst DNA stain. Scale bar, 10 µm. K) Example FACS plots of WT or lamin A R377I-expressing epicardial cells treated with vehicle or 5 µM MG132 for 18 hours, then stained for LMNA/C. L) MFI quantification (mean and SD) of 4 independent experiments exemplified by (K). \*\**P* = 0.0038 by two-tailed Welch’s *t*-test.

To test the ability of Lamin A variants to replace endogenous Lamin A/C, we knocked out the endogenous *LMNA* locus in SLICE hiPSCs (Fig. 1 supplement 2A). *LMNA* is very lowly expressed and dispensable in pluripotent cells^16,17^ and *LMNA* KO SLICE hiPSCs retained normal expression of pluripotency markers and exhibited normal stem cell morphology (Fig. 1 supplement 3A-B). In contrast, *LMNA* is highly expressed and functionally important in many differentiated lineages, including cardiac cell types^18^. *LMNA* KO SLICE hiPSCs can differentiate into proliferative epicardial cells (epiCs)^19,20^ and post-mitotic ventricular cardiomyocytes (VCMs)^21^ (Fig. 1 supplement 2D, Fig. 1 supplement 3A-B), albeit with moderately decreased efficiency that became apparent in VCM competition differentiations performed with equal numbers of WT and *LMNA* KO hiPSCs (Fig. 1 supplement 3C,D). *LMNA* KO epiCs and VCMs exhibited abnormal nuclear morphology and displacement of the Lamin A/C-binding protein emerin (Fig. 1 supplement 2D-E), consistent with Lamin A/C loss of function. Reintroduction of WT Lamin A into the SLICE landing pad caused constitutive overexpression of Lamin A in hiPSCs (Fig. 1 supplement 2A-B) with negligible effects on expression of pluripotency markers (Fig. 1 supplement 3B). SLICE-mediated Lamin A expression was maintained over differentiation (Fig. 1 supplement 2B-C). Nuclear morphology and emerin localization were improved by Lamin A re-expression in both lineages (Fig. 1 supplement 2D-E), indicating that SLICE-mediated expression of Lamin A approximates the functions of endogenous Lamin A/C.

We next replaced endogenous Lamin A/C with a library of human Lamin A coding sequence variants generated by <u>d</u>eep <u>i</u>ndel <u>m</u>issense <u>p</u>rogrammable <u>l</u>ibrary <u>e</u>ngineering (DIMPLE)^22^. This ∼18,000 variant library includes missense, nonsense, and synonymous substitutions, as well as in-frame insertions and deletions (indels) of one, two, or three amino acids at each position (Fig. 1A). SLICE integration and selection yielded ∼95% coverage of the variant library in the landing pad (Fig. 1B, Fig. 1 supplement 1E), thus enabling saturation-scale variant library screens in hiPSCs and their differentiated progeny.

### Establishing Lamin A protein abundance screens in hiPSCs and cardiac lineages

To comprehensively profile the effects of sequence variation on Lamin A abundance, we conducted protein abundance screens in SLICE Lamin A library hiPSCs, epiCs, and VCMs by immunostaining and FACS (Fig. 1C, Fig. 1 supplement 1F). We expressed Lamin A variants without an epitope tag, as tagging disrupts key properties of the lamina^23,24^ and may promote aggregation^25^. Instead, we used validated antibodies to detect untagged variants coupled with gentle zinc fixation to facilitate genomic DNA extraction. Fixed and permeabilized hiPSCs, VCMs, and epiCs were immunostained with a monoclonal antibody raised against residues 2-29 of human Lamin A/C, then sorted into equal quartiles of fluorescence intensity by FACS (Fig. 1C, Fig. 1 supplement 4A). Genomic DNA was extracted from each sample, followed by amplification from the landing pad, deep sequencing, and quantification of variant enrichment across quartiles using Lilace, a Bayesian statistical model developed for FACS-based DMS screens^26^ (supplementary table 1). We verified that synonymous variants uniformly clustered around zero abundance effect, while nonsense mutations decreased protein abundance (Fig. 1 supplement 4G). Mutations to residues 1-6 of the N-terminal antibody epitope completely ablated protein detection (Fig. 1D), while clusters of mutations elsewhere in the protein sequence also decreased signal (Fig. 1D-E). To distinguish between protein abundance and epitope masking effects, we conducted a secondary screen using an antibody that recognizes an epitope corresponding to residues 598-611 in the Lamin A tail (Fig. 1 supplement 4A, supplementary table 2) and confirmed a similar pattern of signal loss while mapping this antibody epitope to residues 602-610 (Fig. 1 supplement 4B). The concordance between antibody screens indicates that signal loss reflects decreased protein abundance (Fig. 1 supplement 4C-D). Altogether, these datasets provide the first comprehensive analysis of the effects of Lamin A sequence variation on protein abundance in pluripotent and disease-relevant cardiac cell types.

### Indel and missense mutations destabilize folded domains of Lamin A

Lamin A mutations could decrease protein abundance by inducing degradation or increase abundance due to aggregation. However, our screens indicated that very few mutations significantly elevated protein abundance (Fig. 1D-E). The aggregation propensity of one pathogenic mutation to *LMNA*, the HGPS-causative mutation, has been extensively documented, making it an ideal positive control for our platform^7,27,28^. As this variant is a splice isoform of Lamin A and was thus not included in our library, we introduced it into *LMNA* KO hiPSCs by SLICE to evaluate its effects on protein localization and abundance. Progerin formed perinuclear aggregates in hiPSCs (Fig. 1 supplement 5A) and accumulated to higher levels than WT Lamin A in VCMs (Fig. 1 supplement 5B), confirming the aggregation and accumulation propensity of this toxic GOF mutant in our system. In contrast, microscopy analysis of DMS library-expressing hiPSCs, epiCs, and VCMs did not reveal evidence of aggregates as a common outcome of Lamin A mutations (Fig. 1 supplement 5C). Instead, Lamin A variants previously described to be aggregation-prone when tagged and overexpressed^29^ had significantly decreased protein abundance in our screens (Fig. 1 supplement 5D).

We observed discrete regions within the Lamin A sequence that were especially sensitive to mutations across all cell types. To identify factors contributing to mutation sensitivity, we constructed a random forest model including a range of physicochemical parameters. This model predicted strong relationships between protein abundance effect and evolutionary conservation (ConSurf^30^) as well as predicted thermodynamic destabilization (thermoMPNN^31^) (Fig. 1 supplement 4H-I). Overall, highly conserved residues are most destabilized by mutations (Fig. 1F). Of the most conserved residues, those predicted to be buried within the structure were more mutation-sensitive than those predicted to be surface-exposed (Fig. 1G). This relationship was clearly evident within the conserved globular Ig-like domain of Lamin A, which forms a β-sandwich structure through hydrophobic residue contacts at its center^32^. Within this domain, mutation-sensitive positions were predominantly buried within the β-sandwich (Fig. 1H). Moreover, Ig-like fold abundance scores were inversely correlated with FoldX-predicted^33^ thermostability (Fig. 1I), indicating that structural perturbation plays a role in the observed loss of signal in our screens.

These correlations suggest that mutations to sensitive residues result in misfolding and degradation by the pQC machinery. To directly test this prediction, we reintroduced WT or mutant Lamin A cDNAs into the *LMNA* KO SLICE landing pad with controlled, single-copy expression. This enabled us to interpret a steady state abundance difference as protein-level regulation. Reintroduction of a variant with a very low abundance score, R377I, into epiCs revealed very low steady-state abundance and strong accumulation upon proteasomal inhibition (MG132, Fig. 1J-L), indicating that low steady-state abundance in our screens reflects high rates of protein turnover.

### Relationships between pathogenicity and protein stability

Premature lamin truncations cause LOF and are frequently pathogenic^4^. We hypothesized that destabilizing missense mutations could have similar effects and explored the relationship between missense variant abundance and pathogenicity. Lamin A abundance correlated well with AlphaMissense-predicted pathogenicity^34^ (Fig. 2A). To extend this finding, we evaluated the frequency of low-abundance Lamin A variants in human populations. We defined missense variants at or below the 5^th^ percentile of the VCM synonymous variant distribution as “lowly expressed”, then cross-referenced these to the gnomAD^35^ and Regeneron Million Exomes^36^ databases. These lowly expressed variants were significantly underrepresented in the human population compared to randomly sampled variants matched for mutation type composition, consistent with selection against destabilizing mutations (Fig. 2B). Further, protein-altering variants annotated in the ClinVar database as pathogenic (P) or likely pathogenic (LP) had lower abundance than those annotated as benign (B) or likely benign (LB), supporting a relationship between protein destabilization and human pathogenicity (Fig. 2C). To specifically evaluate cardiac pathology, we measured the incidence of *LMNA*-mediated cardiovascular disease reported in the All of Us database^37^ for individuals harboring lowly expressed *LMNA* variants. Strikingly, these individuals had significantly higher rates of heart failure, atrial fibrillation, AV block, and ventricular arrhythmia than individuals carrying normal-abundance variants or the overall population.(Fig. 2D). This serves as population validation of the pathophysiologic impact of variants on protein stability. To assess the strength of this relationship, we computed OddsPath ratios following the ClinGen Sequence Variant Interpretation framework^38^. Because benign missense variants are rarely reported in ClinVar, we selected 43 additional common missense variants (minimum of twice the maximum allele frequency of pathogenic variants) from the GnomAD and Regeneron exome sequencing databases as a benign proxy (supplementary table 3). For variants with abundance scores less than 12.5% of the synonymous variants, the OddsPath ratio was 5.24, indicating that low abundance in our assay constitutes moderate evidence of pathogenicity.

**Figure 2.**
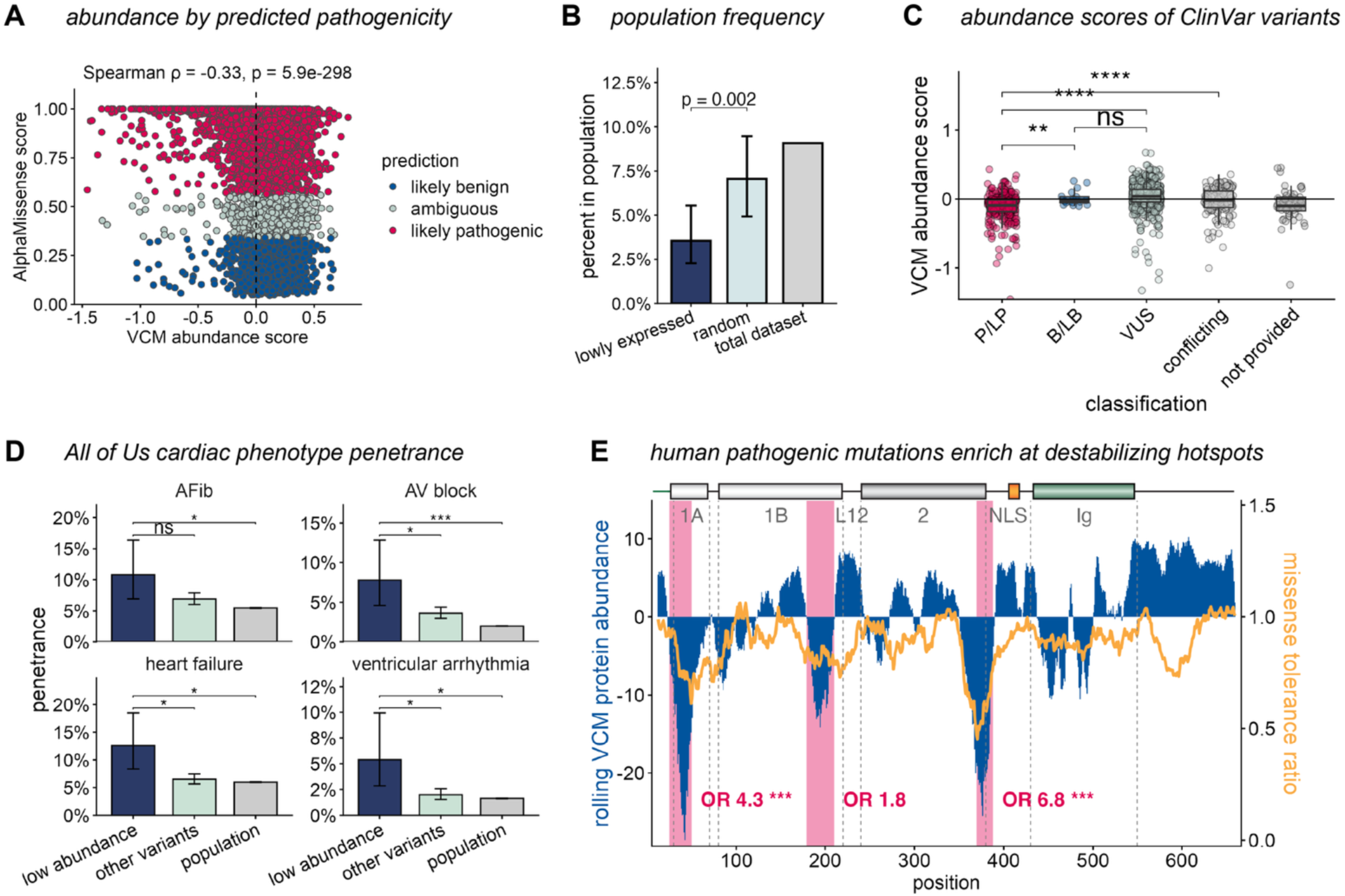
Low lamin A protein abundance is associated with pathogenicity. A) Spearman correlation between VCM variant abundance score and AlphaMissense predicted pathogenicity, colored by prediction category. B) Percentage of variants found in the combined gnomAD and Regeneron population databases (*n* = 1,462 unique variants). Bars: lowly expressed variants (dark blue, VCM abundance < 5th percentile of synonymous scores and Lilace FDR < 0.05), mutation-type–matched random permutations (light blue), and the total dataset (grey). For the lowly expressed set, error bars denote the 95% Wilson confidence interval for the observed proportion; for the random set, error bars denote 2.5th–97.5th percentile interval of the null distribution from 1,000 mutation-type–matched permutations. *P* value, one-sided permutation *P* value testing whether the lowly expressed set is depleted relative to the random null. Random permutations were sampled to match the nonsense/missense/synonymous composition of the test set. C) VCM protein abundance scores of variants with annotations in ClinVar database. Each variant horizontally jittered over boxplots (center line, median). Grouped by ClinVar classification (P/LP: pathogenic/likely pathogenic; B/LB: benign/likely benign; VUS, variants of uncertain significance). \*\**P* < 0.01; \*\*\*\**P* < 0.0001 by two-sided Wilcoxon rank-sum tests with BH correction. D) Cardiac phenotype penetrance among *LMNA* variant carriers by VCM abundance class. Carriers of low-abundance variants (*n* > 160) are compared with carriers of other variants (*n* > 2,800) and with the general All of Us population. Bars show the percentage of carriers in each group affected by each phenotype; error bars are 95% Wilson confidence intervals. Significance was assessed by Fisher’s exact test with Benjamini–Hochberg correction (ns, not significant; \**P* < 0.05, \*\**P* < 0.01, \*\*\**P* < 0.001). E) VCM lamin A protein abundance compared to pathogenic position windows. Each blue bar is a 10-residue sliding window (1-position step) along lamin A, plotted at the window center (x-axis). Red bars, rod N-terminus (positions 25–50), coil 1b C-terminus (179–188), and rod C-terminus (370–388). OR, Fisher’s exact odds ratio for enrichment of pathogenic variants inside window relative to outside. \*\*\**P* < 0.001 by Fisher’s exact test. Yellow line, Regeneron Million Exomes database missense tolerance ratio, where low values indicate fewer alleles present in the population than expected in each rolling window.

Across all cell types, Lamin A was most severely destabilized by mutations to three short segments of the central rod domain (Fig. 1D). To determine if these hotspots relate to human pathogenicity, we tested whether known pathogenic variants were enriched in these windows relative to the rest of the protein sequence. Pathogenic variants were 4.3- and 6.8-fold more likely than expected to fall in coil 1a and the end of coil 2, respectively; the coil 1b window showed the same trend but did not reach significance (Fisher’s exact test; Fig. 2E). This pattern mirrored the missense tolerance ratio (MTR) calculated from the Regeneron Million Exomes dataset^36^, which shows that missense variants in these windows appear in the population less often than expected. Altogether, these results demonstrate that protein destabilization, variant pathogenicity, and low population frequency converge on specific hotspots within the Lamin A rod domain.

### Insights into the structure of the lamin A unit dimer

We hypothesized that mutation-sensitive regions mark surfaces important for Lamin A folding and/or assembly. To explore this possibility, we mapped our abundance data to known and predicted structures of the lamin filament. The domain with the most potently destabilizing mutations was the central rod domain, which mediates the polymerization of the lamins and is responsible for their mechanical properties. This domain is comprised of homodimeric coiled-coil helices separated by linker sequences. Coiled-coil dimers arise from heptad repeats of hydrophobic and charged amino acids, where hydrophobic residues at heptad positions *a* and *d* interact between two alpha helices, flanked by charged residues at *e* and *g* positions that stabilize the twisting filament (Fig. 3A). Our library included in-frame insertions of G, GS, and GSG, as well as deletions of one, two, and three residues at each position. As such indel mutations severely disrupt protein folds^22^, we postulated that they would disrupt the heptad repeat phasing of the rod domain’s coiled coils. In support of this prediction, aligning mean indel abundance scores across the lamin sequence revealed that indels within annotated coiled-coils were more destabilizing than those in linkers (Fig. 3 supplement 1A-B). This reduced sensitivity was especially evident in the L1 (residues 71-80) and L12 (residues 219-242) linkers (Fig. 3 supplement 1A-B), consistent with these segments being flexible in structure^39–41^. Interestingly, we also observed decreased indel sensitivity at a putative linker early in coil 2 (residues 278-288) and near a stutter in the coil 2 heptad repeat (residues 327-337) (Fig. 3 supplement 1C). Altogether, these observations imply that the rod domain contains stable coiled-coil segments punctuated by flexible linkers that could allow for the compression and sliding of lamin filaments, as previously proposed^39,40^. Further, these data indicate that disruption of the coiled-coil rod domain is a major trigger of lamin destabilization.

**Figure 3.**
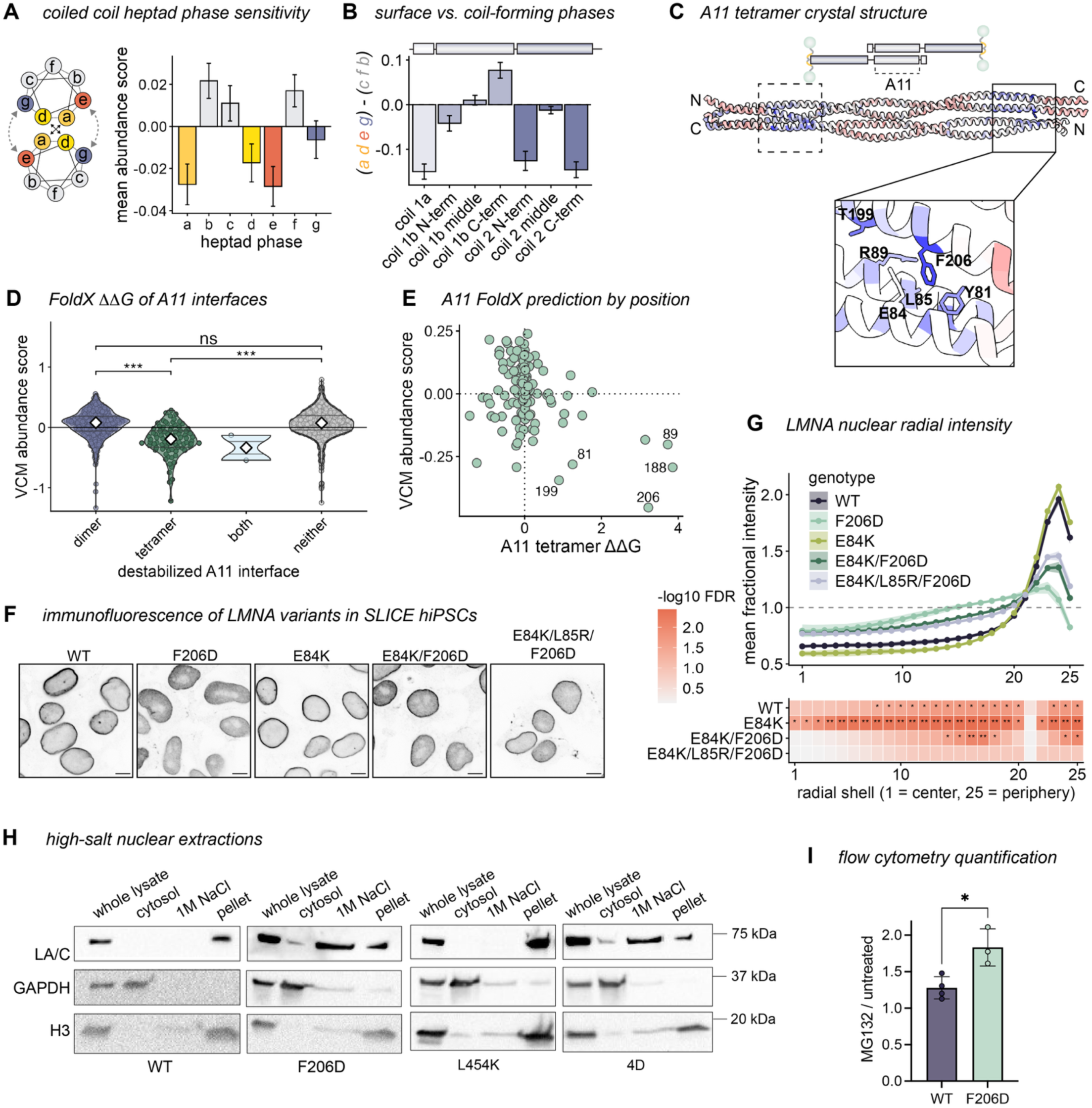
Disrupting lateral polymer assembly destabilizes lamin A. A) Left: coiled-coil diagram showing core *a* and *d* heptad phases interacting across coiled-coil dimer, flanked by charged *e* and *g* positions. *b*, *c*, and *f* positions are surface-exposed. Right: bar plot of mean VCM protein abundance ± standard deviation for each heptad phase within the rod domain, colored by heptad phase. B) Rod domain sensitivity to coiled-coil-forming heptad phases (*a*, *d*, *e*, *g*) versus surface-exposed heptad phases (*b*, *c*, *f*). Bars indicate the difference in mean scores between the two phase groups across lamin subdomains (coil 1a: positions 28–70, linker 1: 71–80, coil 1b N-term: 81–115, coil 1b middle: 116–170, coil 1b C-term: 171–218, linker 12: 219–242, coil 2 N-term: 243–280, coil 2 middle: 281–349, coil 2 C-term: 350–388). C) A11 tetramer crystal structure (PDB 6JLB) colored by positional mean VCM lamin A protein abundance. Boxes: conserved “knob/pocket” interfaces stabilize the tetramer. Antiparallel interface indicated by dashed line. Zoomed interface colored by lamin A abundance and labeled with constituent residues. D) VCM abundance scores for variants predicted by FoldX mutational scan to destabilize the A11 tetramer (PDB 6JLB), either constituent dimer, both, or neither interface. Each variant jittered under violin plots (center diamond, median). Destabilization defined as interaction ΔΔG > 1.6 kcal/mol. \*\*\**P* < 0.001 by pairwise Wilcoxon rank-sum test with FDR adjustment. E) Scatter plot of FoldX predicted A11 tetramer interaction ΔΔG versus positional mean abundance scores, with destabilizing residues labeled. F) Immunofluorescence images of *LMNA* KO cells expressing indicated lamin A single, double, and triple mutants in SLICE landing pad, stained with N-terminal LMNA/C antibody. Scale bar, 10 µm. G) Lamin A radial distribution. Fractional intensity (shell mean ÷ whole-nucleus mean) from nuclear center (shell 1) to periphery (shell 25); mean ± SEM across 3 biological replicates (*n* = 120–172 nuclei per genotype). Lower strip: per-shell paired *t*-tests versus F206D (paired by replicate), BH-FDR-corrected across shells; tile color = −log₁₀(FDR), ** < 0.01, * < 0.05. H) Western blots of high-salt nuclear fractionations. *LMNA* KO hiPSCs expressing the indicated variants from the SLICE landing pad were snap frozen and gently lysed by freeze/thaw. Nuclei were extracted with 1 M NaCl and separated into soluble and insoluble fractions, then blotted for LMNA/C, GAPDH (cytosolic marker) and histone H3 (nuclear marker). I) MFI quantification (mean and SD) of 3 independent experiments represented by (Figure 3 supplement 2C). \**P* = 0.0427 by two-tailed Welch’s *t*-test.

We noted that the effect of missense mutations to the coiled coils depended on their position in the tertiary structure. This pattern of sensitivity followed heptad repeat phasing (Fig. 3A, Fig. 3 supplement 1D) and was strongest near the termini of most coiled coil segments (Fig. 3B, Fig. 3 supplement 1B). Heptad repeat hydrophobic core (*a* and *d*) and electrostatic flanking positions (*e* and *g*) were generally enriched for destabilizing mutations, indicating that mutations that interfere with the coiled coil dimer destabilize Lamin A (Fig. 3A, Fig. 3 supplement 1D). However, the pattern of sensitivity at the C-terminal part of coil 1b was reversed compared to the other coils. There, outward-facing *f* and *c* positions were most destabilized by mutations, indicating that extradimeric interactions at this interface facilitate Lamin A protein stability (Fig. 3B).

### Lateral tetramerization via the “A11” interface contributes to Lamin A stability

Coil 1b of the Lamin A rod domain constitutes a higher-order assembly interface universal to all IFs, known as the “A11” interface^42^. The lamins polymerize laterally and head-to-tail to form a vast network of 3.5 nm-wide filaments^43^. The lateral A11 tetramer has been captured by X-ray crystallography^44^; here, conserved residues on the outer surface of coil 1b interact between two antiparallel coiled-coil dimers. These conserved residues correspond to the destabilizing outward-facing heptad positions we identified in our screens (Fig. 3B-C). To probe the extent of the relationship between Lamin A destabilization and disruption of the A11 interface, we used FoldX^33^ to predict the consequences of mutations to this interface on tetramer thermostability. Mutations predicted to disrupt the tetramer had lower protein abundance scores than those predicted to disrupt the coiled-coil dimers (Fig. 3D, Fig. 3 supplement 2A). The surface-exposed positions 188, 199, and 206, as well as their antiparallel binding partners 81 and 89, had the highest predicted ΔΔG values and the lowest protein abundance scores (Fig. 3E).

To investigate the effects of mutations to the A11 interface, we reintroduced one strongly destabilized variant, F206D, into *LMNA* KO hiPSCs via SLICE. In the A11 crystal structure, F206 occupies an outward-facing heptad position *c*, where it bridges the interaction with a second coiled coil dimer at L85 and Y81 (Fig. 3C)—a conserved, hydrophobic “knob/pocket” contact shared among IFs^42^. Immunofluorescence microscopy revealed that the F206D mutant did not incorporate into the lamina and was instead mislocalized to the nuclear interior (Fig. 3F-G). To test whether the mislocalization of F206D was in fact caused by disruption of the A11 tetramer, we made compensatory charge-neutralizing mutations on the other surface of the A11 interface (E84K and L85R). Both the F206D/E84K and F206D/E84K/L85R mutants partially rescued peripheral enrichment (Fig. 3F-G), consistent with disruption of the A11 multimer interface, rather than disruption of the unit dimer, underlying F206D mislocalization.

Displacement from the nuclear lamina is indicative of Lamin depolymerization. To assess the assembly status of our mutants, we compared them to a known assembly-defective mutant, with phosphomimetic substitutions at four key mitotic phosphosites (T19D, S22D, S390D, S392D; termed 4D). As phosphorylation disassembles the nuclear lamina shortly before mitosis^45^, this mutant benchmarks the localization of a disassembled lamin protein. Surprisingly, the F206D variant was displaced from the nuclear periphery even more dramatically than the SLICE-expressed 4D phosphomimetic mutant (Fig. 3 supplement 2A-B). As assembled lamins are highly insoluble, we also measured assembly of Lamin A variants by biochemical fractionation. These experiments revealed that both F206D and the 4D mutant were more readily solubilized in high salt buffer than wild-type Lamin A (Fig. 3H), consistent with a failure of these mutants to incorporate into the insoluble lamina meshwork. The F206D variant also accumulated more in epiCs than WT Lamin A upon proteasome inhibition with MG132 (Fig. 3I, Fig. 3 supplement 2C), indicating that this mis-assembled variant is recognized and degraded by the proteasome.

To understand whether these effects were unique to A11 interface mutations, we evaluated the consequences of a destabilizing mutation to the Ig-like domain, which has no known role in lamin polymerization, on lamina assembly. While this mutation (L454K) destabilized Lamin A, it remained more enriched at the nuclear periphery than F206D (Fig. 3 supplement 2A-B) and remained insoluble after high-salt extraction (Fig. 3H), indicating that it incorporates normally into the lamina meshwork. Together, these results indicate that distinct, domain-dependent mechanisms control the stability of the Lamin A protein: mutations at the A11 lateral assembly interface both depolymerize and destabilize the Lamin A protein, whereas mutations to the Ig-like domain reduce protein abundance without altering incorporation into the lamina meshwork.

### A mutation-informed model of the Lamin A head-to-tail polymerization interface

IFs polymerize longitudinally by head-to-tail association between the N- and C-termini of adjacent rod domains, forming the “ACN” assembly interface^42^. Unlike the A11 interface, no high-resolution structure of the ACN interface exists. Further, the degree and conformation of the ACN overlap is contested; previous crosslinking studies have suggested the coils interact either by intercalation of the coiled coil dimer ends^46^ or by lateral association via their adjacent unstructured domains^40^. Because our abundance screen sensitively highlighted A11-forming residues, we looked for similar patterns at the rod domain termini to understand the nature of Lamin A’s ACN assembly.

The ends of Lamin A’s rod domain were the most mutation-sensitive regions in our fitness and abundance screens (Fig. 4A, Fig. 1D-E), leading us to predict that these residues are involved in ACN assembly. Consistent with this idea, we noted a remarkably symmetrical pattern in their abundance scores: both termini contained a ∼25 residue stretch of sensitivity, flanked by regions of relative mutational tolerance (Fig. 4A). To gain structural insight into this putative interface, we modeled the interaction between the N-terminus of one coiled coil dimer with the C-terminus of another using AlphaFold3^47^ (Fig. 4B-C; Fig. 4 supplement 1A-B). The resulting structural model (ipTM = 0.61, pTM = 0.63) predicted the intercalation of the two dimers with an overlap of 4.2 nm (Fig. 4C)—closely matching the dimensions of the destabilized regions in our screen. Residues at the most mutation-sensitive positions formed confidently aligned contacts along the tetramer interface (min. predicted aligned error 2.5-3.5 Å; Fig. 4 supplement 1A). To test whether defects in this modeled interface reduce protein abundance, we again used FoldX^33^ to predict the energetic effects of mutations on the modeled ACN tetramer versus each constituent dimer. This analysis indicated that variants predicted to destabilize only the ACN tetramer, or both the tetramer and the unit dimer, had lower abundance scores than those predicted to affect the unit dimer only (Fig. 4D), implying that ACN assembly defects reduce protein abundance in cells. As a complementary approach to identify potential ACN contacts, we searched for evidence of evolutionary covariation at Lamin A’s N and C termini using EVcouplings analysis^48^ (Fig. 4E). Notably, this analysis identified significant evolutionary couplings between the C-terminal residue Y376 and the N-terminal residue N39 (probability = 0.73, score = 1.02) and between Y376 and L38 (probability 0.57, score 0.28), both of which lie in the heart of the AlphaFold3 model and were strongly sensitive to mutations in our screens (Fig. 4C). Finally, the AlphaFold3 model predicted that coil 1a residues at heptad position *e* aligned with coil 2 residues at position *g* (Fig. 4 supplement 1C), indicating rearrangement of coiled coil electrostatic interactions to accommodate coil intercalation. Strikingly, this alignment was reflected in mirror-image sensitivity patterns in our screen data: the most strongly destabilizing mutations were found at *e* positions in coil 1a and at *g* positions at the end of coil 2 (Fig. 4 supplement 1D).

**Figure 4.**
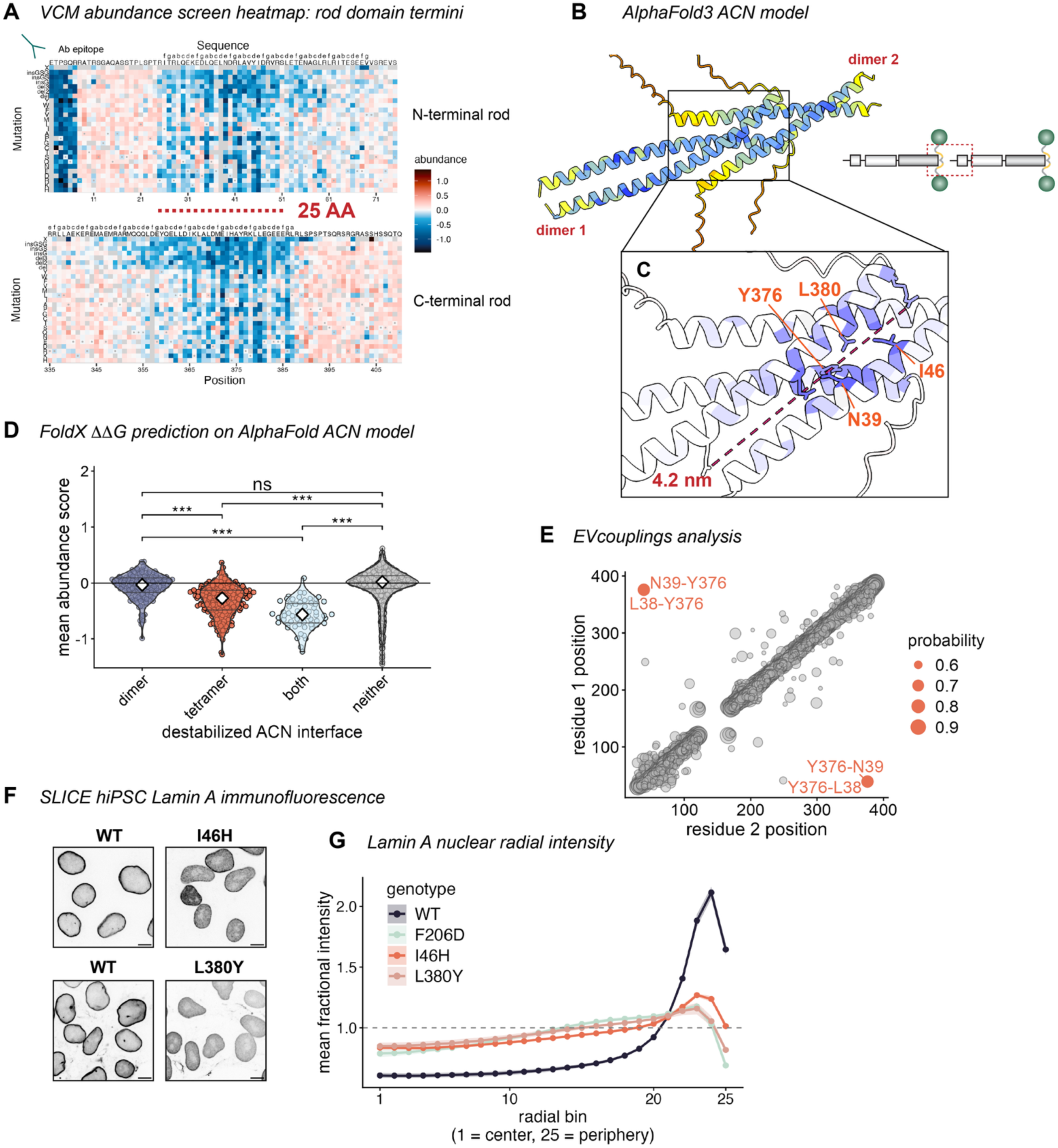
Destabilizing mutations highlight head-to-tail multimerization interface. A) Rod domain terminal heatmaps (top: positions 1–75, bottom: 335–410) colored by lamin A protein abundance in VCMs. Position numbers annotated along the bottom, WT amino acid and heptad phase along the top, and mutant amino acid on the left. N-terminal antibody epitope indicated. B) AlphaFold3 ACN model colored by predicted local Distance Difference Test (pLDDT), a per-residue confidence measure. N- and C-terminal coiled-coil dimers indicated in red. C) Zoomed interface view of AlphaFold3 ACN model, colored by positional mean VCM protein abundance with low-abundance residues labeled. Dashed line indicates overlap length between N- and C-terminal coils. D) VCM lamin A protein abundance scores for variants predicted by FoldX mutational scan to destabilize the AlphaFold3 ACN model tetramer, either constituent dimer, both, or neither interface. Each variant jittered under violin plots (center diamond, median). Destabilization defined as interaction ΔΔG > 1.6 kcal/mol. \*\*\**P* < 0.001 by pairwise Wilcoxon rank-sum test with FDR adjustment. E) EVcouplings analysis of lamin A rod domain. Each point is a confident co-evolving residue pair (EC score > 0.2) plotted at its two sequence positions symmetrically about the diagonal. Point size: EVcouplings coupling probability. The N39–Y376 pair is highlighted in orange and obscures the smaller L38–Y376 pair. Their distance from the diagonal reflects an unusually large sequence separation for two strongly coupled residues, consistent with a direct contact across the ACN interface. F) Immunofluorescence images of *LMNA* KO cells expressing indicated lamin A mutants in SLICE landing pad, stained with Ig-fold LMNA/C antibody. Scale bars, 10 µm. G) Fractional intensity (shell mean ÷ whole-nucleus mean) from nuclear center (shell 1) to periphery (shell 25). *n* = 216–364 nuclei per mutant across independent imaging experiments (WT, *n* = 11; F206D, *n* = 4; I46H, *n* = 3; L380Y, *n* = 3). Lines are means across biological replicates, ribbons, SEM across replicates. WT-normalized I46H and L380Y means were compared to F206D by Welch’s unpaired *t*-test, and no shell was found to be significantly different.

To test these predictions of ACN engagement, we examined variants at sensitive and conserved hotspot positions—I46H, N39I, Y376Q, and L380Y—by SLICE-mediated reintroduction into *LMNA* KO hiPSCs. Each of these variants was displaced from the nuclear periphery and was more soluble by high salt extraction, indicating impaired assembly into the lamina meshwork (Fig. 4F-G; Fig. 4 supplement 1E-F). These outcomes are consistent with depolymerization and similar to the effect of the A11 interface mutant F206D (Fig. 3F-H). Altogether, these results converge on a model of the Lamin A ACN interface that involves interdigitation of the termini of coil 1a and coil 2.

### Consequences of Lamin A variants on cardiomyocyte fitness

Given the strong sensitivity of cardiomyocytes to *LMNA* mutations^4^ we hypothesized that some Lamin A mutations would decrease the fitness of iPS-derived VCMs. To test this, we compared variant representation between VCM and epiC libraries (Fig. 5A). Since both libraries arose from the same hiPSC starting pool, we reasoned that variants that generally interfere with cell fitness or differentiation would be equally depleted from both, while variants specifically deleterious to VCMs would be selectively depleted in this laminopathy-vulnerable cell type. To test for VCM-specific variant toxicity, we computed a log ratio of unsorted VCM to unsorted epiC variant counts using Rosace, a Bayesian statistical model developed for fitness screens^49^. This ratio represents a VCM “fitness score,” with negative values corresponding to variants that are underrepresented in VCMs (supplementary table 4). Nonsense, missense, and indel variants had significantly lower fitness scores than synonymous variant controls, indicating selective depletion of deleterious variants in VCMs (Fig. 5B). Evolutionarily conserved positions had a higher proportion of low-fitness variants than non-conserved positions (Fig. 5 supplement 1A). To classify scores, we applied a confidence interval-based stratification^50^. We calculated 95% confidence intervals for each variant (score +/- 1.96 x SD), which we used to compare against the 5^th^ and 95^th^ percentiles of the synonymous distribution. Cardiac hiPSC differentiation is intrinsically variable. Nevertheless, when we compared the distribution of low fitness variants (<5^th^ percentile of synonymous distribution) across ClinVar classifications, we found that both pathogenic variants and VUS were enriched in low fitness variants relative to benign variants; pathogenic variants were significantly enriched for variants that were both low-fitness and low-abundance, while VUS were enriched for variants with low fitness only (Fig. 5C). Positional mean protein abundance correlated weakly but significantly with VCM fitness (Spearman’s rho = 0.11, *P =* 0.0046) (Fig. 5D; Fig. 5 supplement 1B). This correlation was driven by positions with known P/LP variants, while there was no significant correlation for positions with only documented VUS (Fig. 5D). These observations indicated a small but significant selection effect during VCM differentiation, prompting us to stringently filter the data to uncover residues important for Lamin A function in VCMs.

**Figure 5.**
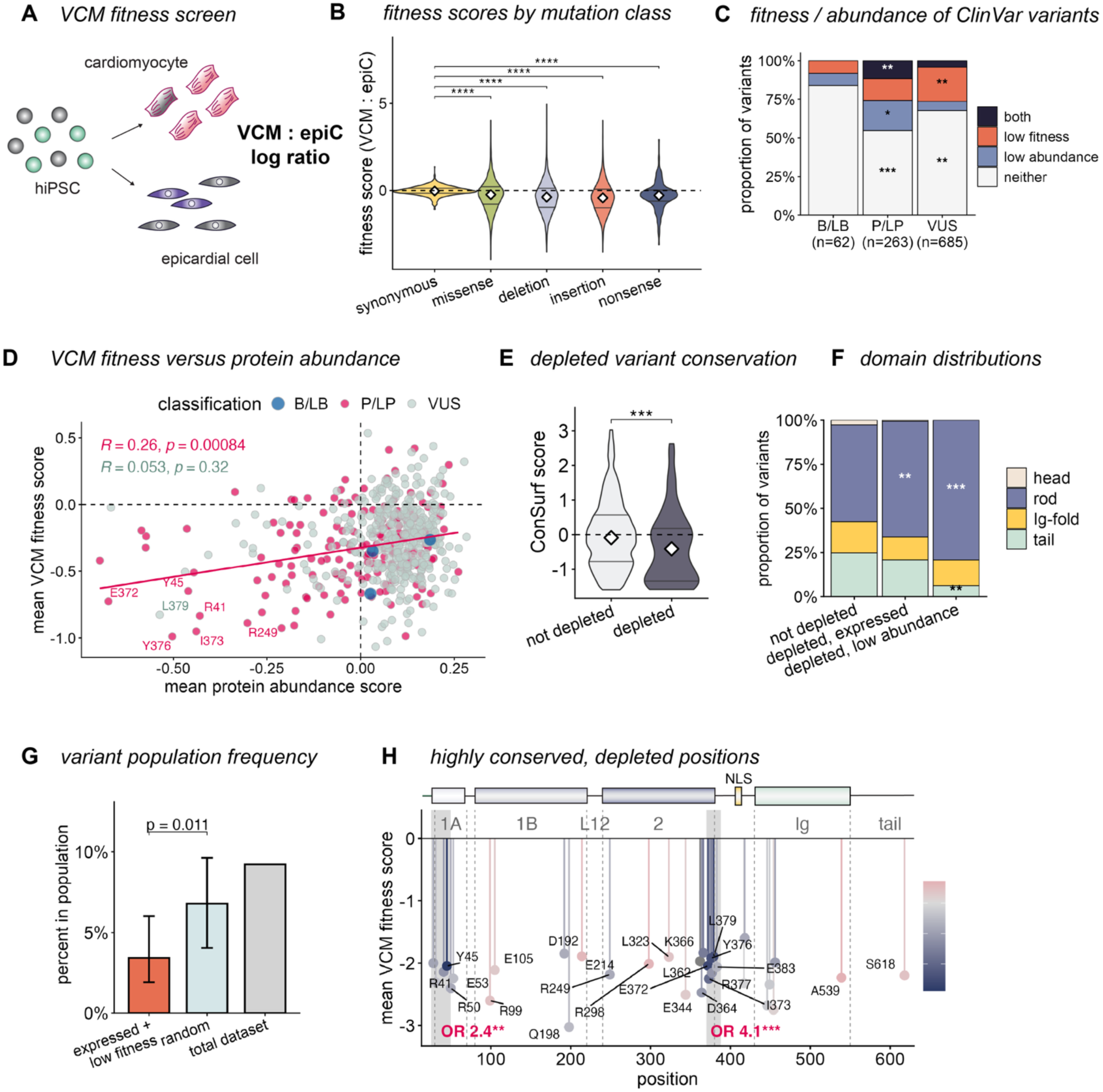
Variant effects on VCM fitness. A) Fitness screen diagram. Variant counts are compared between unsorted VCM and epicardial samples to assess representation after differentiation from the same starting hiPSC pool. Screen tests for underrepresentation of toxic variants (illustrated by grey cells) in VCM pool. B) Distribution of fitness scores by variant type. Violin plots of per-variant Rosace-calculated fitness scores (VCM relative to epiC pools; more negative = greater depletion) for each mutation class: synonymous (*n* = 629), missense (*n* = 11,308), deletion (*n* = 1,647), insertion (*n* = 1,594), and nonsense (*n* = 341). White diamonds: median. Inner lines: 25th and 75th percentiles. \*\*\*\**P*-adj < 0.0001 by pairwise Wilcoxon rank-sum tests against synonymous with Benjamini–Hochberg correction. C) Proportion of ClinVar variants in each clinical category (B/LB, VUS, P/LP) assigned to a combined-screen class: both (low fitness and low abundance), low-fitness only, low-abundance only, or neither. Fitness and abundance hits are variants scoring below the synonymous 5th percentile by Rosace and Lilace analysis, respectively. \**P* < 0.05, \*\**P* < 0.01, \*\*\**P* < 0.001: enrichment or depletion relative to B/LB by Fisher’s exact test. D) Positional mean VCM protein abundance (x-axis) versus fitness (y-axis), colored by most severe ClinVar classification at each position. Red line, linear best fit for P/LP positions. Labeled residues mark low fitness, low abundance positions. Top left, Spearman correlations for P/LP (red) and VUS (grey). E) Violin plot of ConSurf conservation scores for depleted versus not depleted missense variants. More negative ConSurf scores indicate greater evolutionary conservation. White diamonds: median. Inner lines: 25th and 75th percentiles. \*\**P* < 0.01, \*\*\**P* < 0.001 by pairwise Wilcoxon rank-sum test. F) Domain distribution of variants that are not depleted, depleted with low abundance, and depleted but normally expressed. Stacked bars: proportion of each group falling in the head, rod, tail, or Ig-fold. \**P* < 0.05, \*\**P* < 0.01 for enrichment of a region in a depleted group relative to the not-depleted group by Fisher’s exact test. G) Percentage of variants found in the combined gnomAD and Regeneron population databases (*n* = 1,462 unique variants). Bars: low-fitness, expressed variants (see Methods), mutation-type–matched random permutations (light blue), and the total dataset (grey). For the low-fitness set, error bars denote the 95% Wilson confidence interval for the observed proportion; for the random set, error bars denote 2.5th–97.5th percentile interval of the null distribution from 1,000 mutation-type–matched permutations. *P* value, one-sided permutation *P* value testing whether the low-fitness set is depleted relative to the random null. Random permutations were sampled to match the indel/nonsense/missense/synonymous composition of the test set. H) Lollipop plot of per-position mean VCM fitness score for highly conserved (ConSurf score < −0.885), depleted positions along lamin A. Stem height: mean VCM fitness score at each position (more negative = greater depletion); color, mean protein abundance (blue, low; grey, neutral; red, high). Labels mark WT residue and position. Domains are annotated above. Shaded bands: rod domain terminal windows as in (Figure 2E); OR, odds ratio of enrichment of 278 depleted variants within window relative to the rest of the protein. \*\**P* < 0.01, \*\*\**P* < 0.001 by Fisher’s exact test.

We filtered for positions with a high proportion of particularly low-fitness variants (see methods), yielding a list of 292 high-confidence depleted missense variants (supplementary table 5). These variants were significantly more conserved than non-depleted variants (Fig. 5E). While this list included 79 low-abundance variants, 199 had normal protein levels in VCMs, indicating that our fitness data capture effects independent of protein instability. Both normally expressed and low-abundance depleted variants were enriched in the rod domain relative to non-depleted variants (Fig. 5F, Fig. 5 supplement 1C). Further, these depleted but expressed variants were significantly underrepresented in human population databases (Fig. 5G, Fig. 5 supplement 1D), supporting the idea that they could be important for human Lamin A function. Within this normal abundance class, we identified 45 highly conserved residues (supplementary table 5, ConSurf grade 8-9); 36 of these are predicted by ConSurf to be surface-exposed, suggesting they may not induce protein misfolding, but may perturb lamin function by other mechanisms. Together, our fitness and abundance data point to two distinct classes of variants with VCM-specific sensitivity: those that disrupt lamin function without destabilizing the protein, and those that that trigger both protein degradation and VCM toxicity. Interestingly, low-fitness variants were significantly enriched in the destabilizing hotspots at the ends of the rod domain (Fig. 5H; Fig. 5 supplement 1E), recapitulating the accumulation of ClinVar pathogenic variants at these regions (Fig. 2E). P/LP variants with both low fitness and low abundance overwhelmingly concentrated at the rod termini (Fig. 5D), consistent with our prediction that severe destabilizing effects of mutations there contribute to cardiac toxicity.

### Lineage-specific consequences of polymer interface mutations on the lamina

Our fitness screen uncovered VCM-specific sensitivity to the disruption of the rod domain termini. We next asked whether the destabilizing effects of these mutations differed by cell type. To query relative abundance differences between screens, we compared the z-scored positional means of hiPSC, epiC, and VCM abundance scores (Fig. 6A). Interestingly, while mutations to the ACN interface destabilized the protein in all cell types, this effect was more severe in cardiac lineages. N-terminal rod mutations were most severely destabilizing in VCMs, and mutations in the C-terminal rod were more destabilizing in differentiated cells than in hiPSCs (Fig. 6A, Fig. 6 supplement 1A-C). The A11 polymer interface was also more sensitive to mutations in epiCs and VCMs, indicating that these cell types are more sensitive to defects in Lamin A assembly than hiPSCs (Fig. 6A, Fig. 6 supplement 1A-C). We corroborated this observation by measuring the relative steady-state abundance of the A11 interface variant F206D compared to the WT protein after controlled, single-copy reintroduction into *LMNA* KO SLICE hiPSCs and epiCs by quantitative Western blotting. This analysis confirmed that the F206D mutation is intrinsically more unstable in epiCs than in hiPSCs (Fig. 6 supplement 1D-E). In contrast, positions in the Ig-like fold were significantly more destabilizing in stem cells (Fig. 6A, Fig. 6 supplement 1A-C). These findings indicated that differentiated cells were more sensitive to polymer-disrupting mutations, but not to destabilizing mutations in general. Together, our screens reveal that mutations to multimerization interfaces are more potently destabilizing after cardiac differentiation. While the ACN and A11 polymer interface mutations were more destabilizing after differentiation, only ACN interface mutations affected VCM fitness. The consequences of disrupted Lamin A assembly on both protein abundance and function are thus cell type specific.

**Figure 6.**
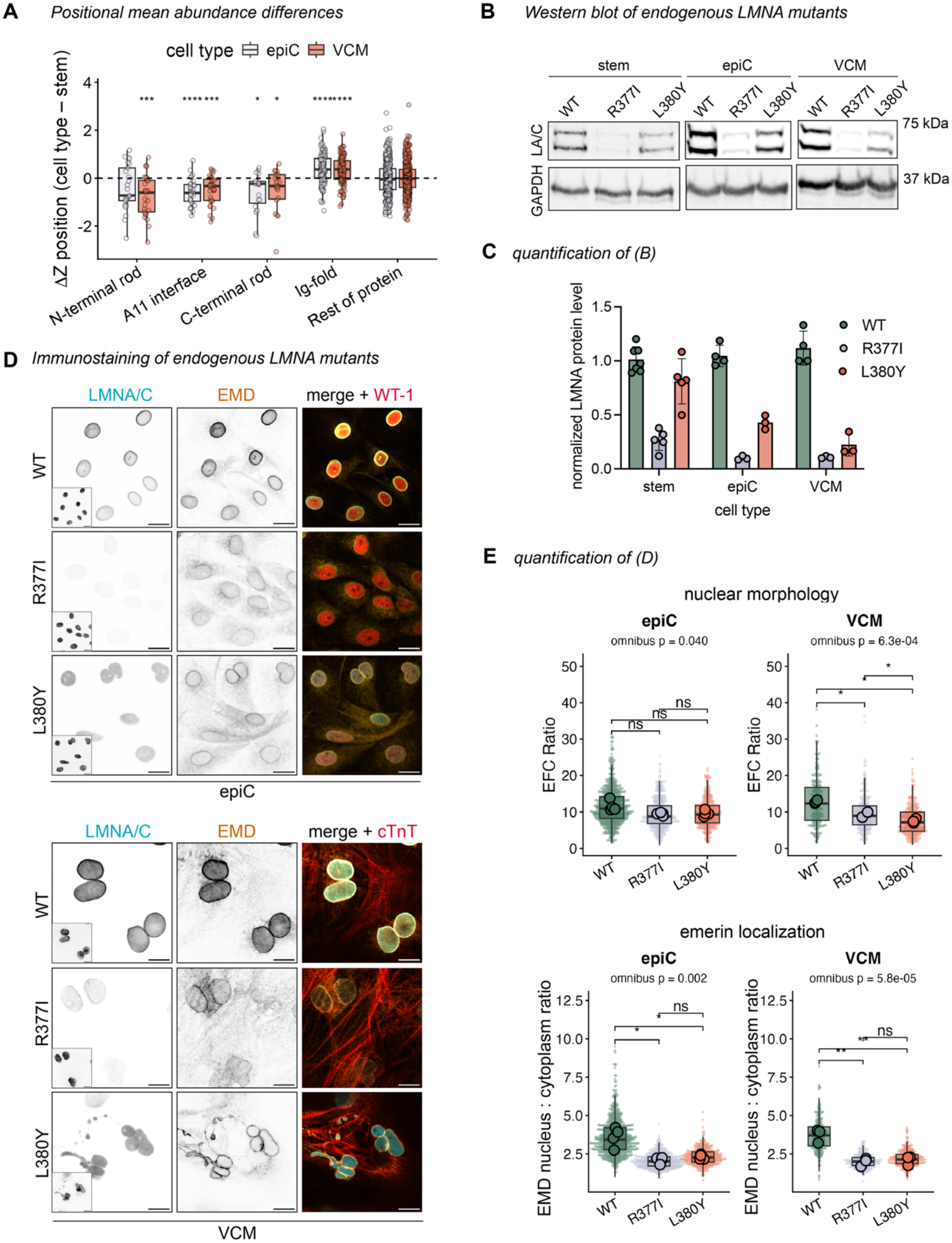
Endogenously expressed ACN variants display cardiomyocyte-specific defects. A) Difference in z-scored positional mean lamin A abundance between differentiated cells (epiC, VCM) and stem cells for the N-terminal rod (positions 25–50), A11 interface (179–210), C-terminal rod (370–388), Ig-fold (430–546), and all other positions. *P* values indicate whether each region’s median differs from 0. \**P*-adj < 0.01, \*\*\**P*-adj < 0.001 by Wilcoxon test with FDR correction. B) Western blot of LMNA/C levels across cell types. hiPSCs CRISPR-edited to express indicated point mutations at endogenous *LMNA* locus were differentiated into epiCs and VCMs, then lysed and blotted for LMNA/C and GAPDH. Representative of 3 independent experiments. C) Quantification of the upper LMNA/C band intensity for Western blots shown in (B). Signal was normalized to mean WT intensity for each cell type. D) Immunofluorescence images of CRISPR-edited differentiated epiCs (left) and VCMs (right), fixed and stained for LMNA/C, emerin (EMD), and lineage markers (epiC: WT1, Wilms tumor 1. VCM: cTnT, cardiac troponin T). Inset, Hoechst DNA stain. Scale bar, 20 µm. E) Quantification of imaging experiments. Box plots show EFC ratio (higher = more regular nuclear morphology, top), emerin nucleus:cytoplasm intensity ratio (bottom). Small points, individual nuclei (box = pooled median and IQR); large points, per-replicate means (the unit of *n*). Omnibus *P*: nested ANOVA (*n* = 4 epiC / 3 VCM biological replicates) indicates difference between all cell types. Brackets: Holm-corrected pairwise paired *t*-tests on replicate means (\**P* < 0.05, \*\**P* < 0.01, \*\*\**P* < 0.001).

To test whether the functional impact of ACN variants depends on endogenous control of protein dose, we introduced homozygous point mutations at two sensitive and conserved C-terminal rod domain positions (L380Y and R377I) into hiPSCs by CRISPR/Cas9 genome editing. We selected these variants based on their severe but distinct phenotypes in our screens: R377I was potently destabilized in all analyzed cell types, while L380Y was most destabilized in differentiated cells (Fig. 6 supplement 1F). L380Y was predicted by FoldX to destabilize the AlphaFold3 ACN model tetramer more potently than R377I (ΔΔG 18.72 versus 2.11 kcal/mol, Fig. 6 supplement 1G), and we noted that these two variants also displayed distinct localization patterns; when expressed from both the endogenous locus (Fig. 6 supplement 2A-B) and the SLICE landing pad (Fig. 4F, Fig. 1J), L380Y localized exclusively to the nuclear interior, while R377I maintained detectable peripheral localization. We differentiated these CRISPR-edited cells alongside control WT clones. Mirroring our screen results, quantitative Western blotting revealed that both endogenously expressed mutant proteins were more potently destabilized in differentiated cells than in stem cells (Fig. 6B-C), while their RNA levels were unaffected (Fig. 6 supplement 2D). The effect of cell type on protein abundance was strongest for L380Y; it was expressed at 81% of WT Lamin A levels in stem cells, 43% in epiCs, and 22% in VCMs (Fig. 6C). These two variants thus typified the pattern of increased instability of polymer-disrupting variants we observed in our abundance screen. We next explored the effects of these endogenously expressed mutants on two lamina-dependent nuclear phenotypes: nuclear shape and nuclear enrichment of a lamina-binding protein. In both epiCs and VCMs, both mutations caused the endogenous Lamin A-binding nuclear membrane protein emerin to mislocalize from the nuclear envelope into the peripheral ER (Fig. 6D-E). In contrast, nuclear morphology was lineage-sensitive: it was unaffected by either mutation in stem cells, marginally affected in epiCs, and strongly affected in VCMs (Fig. 6D-E, Fig. 6 supplement 2A,C). In VCMs, the depolymerized L380Y mutation induced especially severe nuclear deformations, even though this variant was more highly expressed than R377I (Fig. 6B-C). The nuclear morphology defect caused by L380Y mutation was even more severe than that caused by total Lamin A knockout (Fig. 6 supplement 2E), indicating that this effect cannot be attributed solely to decreased lamin expression. Altogether, these results confirm our findings with SLICE-expressed LMNA variants and indicate that the cellular impact of lamin mutations depends on both the molecular consequences to lamin assembly and cell state.

## Discussion

Here, we created and deployed SLICE to conduct the first saturation-scale variant profiling in differentiated cell types, revealing structural features of Lamin A and cell type-specific sensitivity to Lamin A mutations. Our SLICE system leverages recent advances in synthetic biology to efficiently integrate and stably express comprehensive variant libraries in hiPSCs and their differentiated progeny for the first time. This platform can be broadly applied to the controlled analysis of variant effects across cell states; here, we applied deep mutational scanning to Lamin A in pluripotent stem cells, epicardial cells, and ventricular cardiomyocytes.

Our findings provide insight into the dynamic flexibility of Lamin polymers. Indel scanning revealed sensitivity to register shifts within structured domains (the central rod and Ig-like fold), with some surprising exceptions: rod domain linkers were indel-tolerant, even though these appear helical in crystal structures^44^. Moreover, a large portion of coil 2 is tolerant to shifts in heptad register, suggesting a capacity for dynamic conformational change. As the mechanical properties of Lamin A depend on the rod domain^51^, it is tempting to speculate that the flexible regions we identified could play a role in modulating the response to force on the nucleus.

While a complete structure of the lamin filament has evaded the field for many years, our protein abundance screens yielded unique insight into the native lamin polymer. We see a clear pattern of destabilization along the coiled-coil interface at each rod domain terminus. We propose that these surfaces intercalate to form a head-to-tail (ACN) tetramer. Several lines of evidence support this model: first, destabilizing regions at the rod domain termini are symmetrical, demarcated, and align with the AlphaFold3-predicted overlap length of 4.2 nm. Second, we note a significant evolutionary coupling between two strongly sensitive residues at the heart of the predicted interface, Y376 and N39, implying that these residues have co-evolved to preserve their interaction. Third, our abundance scores correlate with predicted destabilization of the AlphaFold3 tetramer and track with the intercalated alignment of the coils by heptad register. Finally, consistent with a role in higher-order assembly, destabilizing variants at the ends of the rod domain are displaced from the nuclear periphery and are more soluble by salt extraction. While no structure of a Lamin ACN has been resolved, coil intercalation or “interlocking” is emerging as a general mechanism of intermediate filament assembly and has been recently proposed to occur in vimentin^52^ and GFAP^53^ filaments.

Our data imply that defects in lateral (A11) and head-to-tail (ACN) Lamin A tetramers are sensed by cellular pQC machineries and trigger protein destabilization. Mislocalized variants at these interfaces (F206D at A11 and R377I at ACN) are poorly expressed but stabilized by proteasome inhibition, indicating that polymerization-defective variants undergo rapid turnover. In contrast to A11 and ACN mutations, destabilizing Ig-like fold mutations such as L454K do not interfere with multimerization but instead perturb subunit folding. Our data thus distinguish at least two routes to lamin turnover: one triggered by defective higher-order assembly and the other by monomer misfolding. Precisely how cells recognize and triage these two classes of variants will require further investigation.

The finding that Lamin A polymer disruption correlates with protein instability is somewhat surprising. Depolymerization is a fundamental part of the lamin life cycle: phosphorylation solubilizes the lamina with every mitosis^45^, and a subset of Lamin A remains depolymerized in the nucleoplasm through interphase^54^. Disassembly alone is therefore unlikely to be sufficient for degradation, and mechanisms must exist to stabilize the soluble pool and prevent its premature turnover. We surmise that polymerization-defective variants expose interaction surfaces that are normally buried within the assembled filament, and that recognition of these exposed surfaces — rather than disassembly per se — marks the protein for degradation. Assembly-coupled pQC has been demonstrated for heterooligomeric protein complexes^55^ and for homodimerizing domains^56^ where pQC factors sense exposed surfaces and route orphan subunits to the proteasome. Interestingly, disruption of type I keratin tetramers also induces protein degradation^57^, hinting that tetramer surface quality control may be an unappreciated feature of intermediate filament biology. Importantly, pQC of Lamin A filaments could shape disease phenotypes, as the spatial and temporal regulation of protein turnover controls both abundance and residual function of misfolded proteins^58^. Identification of these unknown Lamin A pQC machineries may reveal new paths to treatment for laminopathy syndromes.

We demonstrate that mutations to polymer interfaces are more destabilizing in differentiated cardiac cells than in hiPSCs, illustrating the power and importance of interrogating mutation effects in functionally relevant cellular contexts. Endogenously edited mutations to the ACN interface displayed the same lineage-specific patterns of destabilization with the lowest protein abundance in VCMs. However, cardiomyocyte-specific effects of ACN interface mutations may not be due to protein abundance differences alone. While homozygous *LMNA^L380Y/L380Y^* and *LMNA^R377I/R377I^* mutations comparably destabilize Lamin A/C in VCMs, *LMNA^L380Y/L380Y^*induced uniquely severe nuclear deformations in VCMs. These deformations were even more severe than those induced by *LMNA* KO and were not evident in other cell types. As L380Y was completely displaced from the nuclear periphery, we infer that polymer-disrupting mutations are especially detrimental to VCMs. These data urge caution when interpreting reduced protein abundance as “loss-of-function,” as lowly expressed variants may exert toxic effects dependent both upon the functional nature of the mutation and cellular context.

Our Lamin A mutational scans uncovered a convergence between the ACN polymerization interface, pathogenicity, and cardiomyocyte fitness. Mutations to this interface potently destabilize the protein and are toxic to VCMs, indicating a specific sensitivity of cardiomyocytes to polymer-disrupting mutations. This vulnerability is reflected in human genetics, as the ACN interface is significantly enriched for known pathogenic variants. Altogether, these findings imply that some *LMNA* mutations have cardiomyocyte-specific effects. These effects could arise from the convergence of mutations with cardiomyocyte-specific features such as specialization of pQC network composition or activity^6^, protein metabolism^59,60^, or mechanical load.

We did not find protein aggregation or accumulation to be common features of pathogenic *LMNA* mutations. This outcome contrasts with prior studies performed with epitope-tagged, overexpressed variants^25,29^. We infer that Lamin A mutations can cause either decreased protein abundance or aggregation depending on additional factor(s): expression level, epitope tagging, and/or cellular state. We found that unstable mutations are rare in human populations and are frequently linked to cardiac pathogenesis. Our abundance screens meet the “moderate evidence of pathogenicity” standard according to the ClinGen Sequence Variant Interpretation framework^38^, and we identified a subset of strongly destabilized, polymer interface-enriched variants that we predict to be pathogenic (supplementary table 6). Altogether, our fitness and abundance screens highlight functionally distinct classes of variants and detect phenotypes for about half of known pathogenic mutations, indicating that other mechanisms of *LMNA* mutation effects remain to be discovered. In the future, SLICE-enabled deep mutational scanning in differentiated lineages will make it possible to dissect cell type-specific variant effects that drive tissue-restricted disease caused by a variety of mutated proteins.

## Methods

### Cell culture

WTC-11 hiPSCs were obtained from the Berkeley Stem Cell Center. Cells were maintained on Matrigel-coated vessels in mTeSR+ without penicillin/streptomycin. Cells were passaged as clumps (using ReLeSR) for routine culture and as single cells (using Accutase) for counting and electroporating. Cells were plated in ROCK inhibitor Y-27632, then changed to fresh mTeSR+ after 24 hours. Cells were checked for mycoplasma every 3-4 months and were found to be negative.

### Landing pad design and installation

The SLICE landing pad was installed in one allele of the WTC-11 CLYBL safe harbor locus. SLICE relies on the dinucleotide specificity of serine integrases to create directional recognition sites for Pa01 (reef Durrant et al); we placed an attP site with the WT TC dinucleotide on the 5’ end of the landing pad and a mutated CG dinucleotide attP on the 3’ end. A CAGGS promoter-driven BFP-T2A-inducible Caspase 9 (iCasp9) cassette was placed between the recognition sites^61^. Treatment with AP1903 (10 nM, ApexBio) activates the iCasp9, thereby killing cells that retain landing pad expression after donor transfection. CRISPR editing for landing pad installation was performed by electroporating 12.5 pmol pure Cas9-NLS,12.5 pmol sgRNA, and 2 ug HDR template plasmid into 1 million cells using a 100 uL Neon electroporator set to 1150V, 20 ms, and 2 pulses. Landing pad cells were selected in 0.25 ug/mL puromycin for 3 days. BFP+ single clones were FACS sorted into 96-well plates, then expanded and genotyped for heterozygous insertion and absence of plasmid backbone using primers spanning the CLYBL homology arms.

### SLICE donation

SLICE donor constructs were designed to integrate transgenes on the antisense strand of the SLICE landing pad, which we found to be necessary for efficient iCasp9 selection due to promoter interference in semi-integrated cells^62^ (Fig. 1 supplement 1A). To integrate lamin constructs into the SLICE landing pad, we electroporated 1 million cells with 4 ug attB-containing donor plasmid and 1 ug Pa01 integrase plasmid. After 72 hours, we added 10 nM AP-1903 and 10 µM ROCK inhibitor to the media for ∼5 hours, until most cells had lifted. Media was changed to fresh mTeSR+ with AP-1903 and 10 µM ROCK inhibitor overnight, then again the next morning to remove the AP-1903. Cells were expanded and checked by epifluorescence microscopy for pure donor + population. After some SLICE donation/selections, 1-5% of cells did not express the donor. These were sorted away from the donor + population by FACS to obtain 100% donor + cells.

### Lamin A knockout

LMNA knockouts were generated from two landing pad clones. Two sgRNAs were selected to delete ∼300 bp from the first exon, including an endogenous AscI site. Clones were genotyped by PCR and AscI digestion to verify deletion, then by immunofluorescence with N- and C-terminal antibodies to confirm the loss of protein. To verify the sequence of both LMNA alleles, we amplified 2 kb spanning the target edit and sequenced by nanopore long-read sequencing. Raw reads were aligned to the human LMNA sequence with CRISPRnano, which confirmed frameshifting indels on both alleles.

### CRISPR introduction of endogenous missense mutations

Point mutations were introduced into SLICE landing pad WTC-11 hiPSCs by Cas9 ribonucleoprotein (RNP) electroporation with single-stranded oligodeoxynucleotide (ssODN) homology-directed repair (HDR) templates. Guide RNAs (sgRNAs) were selected using the UCSC Genome Browser to position the cut site within 10 bp of the intended edit, then ordered from Synthego. We used the same sgRNA for R377I and L380Y point mutations.

Single stranded, non-complementary ssODN templates were designed (IDT HDR design tool) to incorporate the intended edit together with a silent MluI restriction site insertion and silent ablation of the PAM to prevent re-cutting.

Per reaction, RNPs were assembled by combining 6.4 µg purified Cas9-NLS (Berkeley MacroLab) with 40 pmol sgRNA and incubating 10–20 min at room temperature. hiPSCs were dissociated, counted, and 600,000 cells were resuspended in 9.12 uL Neon genome-editing resuspension buffer. Cell suspension was then combined with the preassembled RNP, 40 pmol ssODN, and 600 ng of a dominant-negative p53 plasmid (p53DD) to ∼12 uL total volume. Cells were electroporated (Neon NXT, 10 µL tip; 1150 V, 20 ms, 2 pulses) and plated onto Matrigel-coated wells in mTeSR+ containing CEPT (Tocris, 7991) and 0.5 µM Alt-R HDR Enhancer V2 (IDT). Medium was changed the following morning and daily thereafter. Cells were maintained in CEPT for 2-3 days after editing to promote survival.

Editing efficiency in the bulk population was assessed by restriction digestion: genomic DNA was extracted, a 747 bp region spanning the edit was PCR-amplified, half the product was digested with MluI (1 h, 37 °C), and digested and undigested products were resolved on agarose gels. Single-cell clones were isolated by limiting dilution into 96-well plates, with CEPT supplementation for the first 2–3 days to support single-cell survival, expanded, and genotyped as above. Positive clones were further sequenced. To resolve alleles, we used primers that spanned a known WTC-11 SNP to amplify the edited region. Only clones homozygous for the edit and heterozygous for the SNP were kept; we found that loss of heterozygosity was common after CRISPR editing, possibly due to large deletions or genomic rearrangement. Clones that received both Cas9 and guide but genotyped as homozygous WT were used as WT CRISPR controls.

### DIMPLE library construction

DIMPLE libraries were cloned as previously described by MacDonald et al^22^. Briefly, the Lamin A coding sequence (UniProt ID P02545) was broken into 15 fragments, each corresponding to a sublibrary of 230 bp oligos. Oligos were designed with BsaI sites flanking encoded mutations (all missense, nonsense, synonymous, insertions of G, GS, GSG, and deletions of 1, 2, and 3 residues). Oligos were synthesized by Twist and were amplified from the pooled library using PrimeStar GXL polymerase and sublibrary-specific primers. PCR reactions were limited to 25 cycles to prevent PCR-induced error propagation. Reactions were cleaned with a Zymo Clean and Concentrate-5 kit. The WT Lamin A ORF was silently mutated to remove BsaI and BsmBI sites and expressed in a Twist High Copy Number kanamycin backbone. For each sublibrary fragment, the plasmid was amplified by PCR using PrimeStar GXL polymerase and sublibrary-specific primers (IDT) to add BsaI sites. 5 uL CutSmart buffer and 1 uL DpnI (NEB) were added to each backbone amplicon and incubated at 37°C for 1 hour, then amplicons were gel extracted using a blue light box and cleaned with a Zymo Gel DNA Recovery kit. 40 uL Golden Gate reactions (200 ng vector amplicon, 20 ng oligo amplicon, 2 uL NEB BsaI-HF V2 Golden Gate Assembly Master Mix, 4 uL 10x T4 ligase buffer) were prepared to ligate oligo sublibaries to their corresponding plasmid backbones. Golden Gate cycling conditions were as follows: (i) 5 min at 37°C, (ii) 5 min at 16°C, (iii) repeat (i) and (ii) 29 times, (iv) 5 minutes at 60°C, (v) hold at 10°C. Reactions were cleaned with a Zymo Clean and Concentrate-5 kit and eluted in 10 uL nuclease-free H2O. Reactions were quantified by NanoDrop and 50 ng were electroporated into 20 uL MegaX DH10B T1^R^ Electrocomp™ Cells (Invitrogen). Cells were recovered without selection for 1 hour in recovery medium, and a small subset was serially diluted and plated to verify that transformation efficiency well exceeded 100x the library size. The remaining outgrowth was cultured in 25 mL LB + 50 ug/mL Kanamycin until the OD reached 0.6. Each sublibrary was purified by miniprep and quantified using a Quant-iT™ dsDNA High-Sensitivity Assay Kit. Sublibraries were pooled in equimolar ratios and then assembled into ampicillin-resistant SLICE donor vectors using BsmBI (75 ng donor vector digested with NotI/XhoI, 75 ng library pool, 1 uL NEB BsmBI-V2 Golden Gate Assembly Mix, 2 uL 10x T4 ligase buffer, 20 uL total). BsmBI Golden Gate cycling conditions were as follows: (i) 5 m in at 42°C, (ii) 10 min at 16°C, (iii) repeat (i) and (ii) 34 times, (iv) 30 minutes at 42°C, (v) 5 min at 60°C, (vi) hold at 10°C. Reactions were cleaned with a Zymo Clean and Concentrate-5 and eluted in 10 uL nuclease free H2O. Final plasmid libraries were electroporated into MegaX DH10B T1^R^ Electrocomp™ Cells, recovered for 1 hour, and spread on 245mm square LB +amp plates. After overnight incubation, the bacterial lawn was scraped from the plate and divided across 3 midiprep columns. Plasmid yield was eluted in nuclease-free H2O and quantified by NanoDrop.

### DMS hiPSC library cell line generation

The DMS library was expressed under the CAGGS promoter (hiPSC and EpiC abundance screens) or the UbC promoter (EpiC and VCM abundance screens). We confirmed that observed abundance differences not an artifact of promoter choice by comparing the two epiC screens to the hiPSC screen (Fig. 1 supplement 4F). The UbC library was electroporated into LMNA KO CLYBL SLICE landing pad cells (37 million cells) as described above for standard SLICE donation. A control electroporation was seeded separately in a 6-well plate; after 3 days, control cells were lifted, counted, and analyzed by flow cytometry to determine integration efficiency. Doubling-time-based calculations (20–24 h doubling) estimated 415,000 surviving cells per million electroporated, and flow analysis showed 1.71% integration efficiency (∼7,062 integration events per electroporation), corresponding to ∼254,000 total integration events and ∼14x library coverage. The main culture was treated with 10 nM AP1903 and ROCK inhibitor 3 days post-electroporation to select for the integrated population, expanded for 4 additional days, and then sorted to purify mCherry+ cells (top 83%, 3.1 million cells). 2 million cells were pelleted for sequencing to assess overall library.

The CAGGS library was introduced into an early version of the CLYBL SLICE landing pad lacking the iCasp9 cassette. Electroporations were staggered across four batches of 15 million cells each and expanded in one T175 flask per batch. Doubling times (20–24 h) were used to estimate post-electroporation survival (1.07–6.8 million cells/batch). Batches were FACS-sorted for mCherry+/BFP-integrants at 3.5–6% efficiency, yielding 105,000–360,000 integration events per sort (∼2.4–20x library coverage), with sort targets scaled to each batch’s estimated coverage. Batches were pooled and sequenced (see below) to assess overall library coverage (∼75% variant representation). Library cells were frozen in 2-million cell aliquots (∼100x library coverage), and sufficient cell numbers were passaged to maintain >100x coverage during routine culture.

### Amplicon sequencing and variant counting

Genomic DNA was isolated using a Zymo quick DNA prep kit, and eluted from the column 3 times with nuclease-free water heated to 70C. The entire gDNA yield was amplified by PCR with Takara GXL polymerase (1000-1500 ng gDNA per 100 uL reaction, 25 cycles of PCR) using landing pad-specific primers, then concentrated with the Zymo clean and concentrate kit. Concentrated samples were separated on a 1% agarose gel and gel extracted with a Zymo gel extraction kit and quantified by Qubit. 1 ng gel-extracted PCR product was library prepped using the Nextera XT prep kit. Samples were sequenced on a NovaSeq X 10B flow cell with paired-end 150-bp reads to a target depth of ∼100×. Raw FASTQ files were processed with the dumpling Snakemake pipeline^49^ (v8.25.5; commit 403e905, accessed 6 December 2024; https://github.com/odcambc/dumpling), which performs QC (FastQC v0.12.1, MultiQC v1.25.2), adapter trimming and mapping with BBTools (BBMap v39.13, Samtools v1.21) and variant calling with GATK AnalyzeSaturationMutagenesis (GATK4 v4.6.1.0), followed by filtering against the designed variant set to generate per-variant counts.

### Epicardial differentiation

WTC-11 hiPSCs were differentiated to epicardial cells via a cardiac progenitor intermediate, following established Wnt-modulation protocols^19,20,63^ with modifications. On day -3, hiPSCs were seeded at 15,000 cells/cm² on Matrigel-coated dishes (Corning 356231) in mTeSR+ with 10 µM Y-27632 (ROCK inhibitor). ROCK inhibitor was withdrawn on day -2 and cells were maintained in mTeSR+ until differentiation. On day 0, cells were rinsed with PBS and switched to cardiac differentiation medium (CDM; RPMI-1640 with 25 mM HEPES and GlutaMAX, 1× B27 minus insulin, and 213 µg/mL L-ascorbate sesquimagnesium salt hydrate (AA2P; Sigma, A8960-5F), containing 10 µM CHIR99021 (Sigma, 361571) for 48 h. On day 2, medium was replaced with CDM alone. On day 3, a 2x solution of IWP-2 (Sigma, 5060720001) in CDM was added directly to the existing medium to a final concentration of 5 µM, and on day 5 medium was replaced with fresh CDM.

On day 7, cardiac progenitors were dissociated with Accutase, gently triturated to a single-cell suspension, and replated 1:5 onto gelatin-coated dishes (Sigma, ES-006) in LaSR replating medium (Advanced DMEM/F12 (Life Technologies, 12634-028) with 2.5 mM GlutaMAX, 146 µg/mL AA2P, 5 µM Y-27632, and 1% FBS). On day 8, medium was changed to LaSR basal medium (Advanced DMEM/F12 with 2.5 mM GlutaMAX and 146 µg/mL AA2P) supplemented with 3 µM CHIR99021. On day 10, cells were returned to LaSR basal medium without small molecules. From the first passage (day 11 or 12) onward, cells were passaged with Accutase (or Versene after the initial passages) at a 1:3–1:5 ratio onto gelatin-coated dishes in LaSR replating medium, and maintained in LaSR basal medium supplemented with 1 µM A83-01 (TGF-β inhibitor, Sigma SML0788) to suppress spontaneous epithelial-to-mesenchymal transition. Epicardial cells exhibited characteristic cobblestone-like morphology and grew in rosette-shaped colonies. Differentiation efficiency was assessed by intracellular flow cytometry (see below) using antibodies against WT-1 (Abcam ab89901, 1:250) and ZO-1 (Proteintech 21773-1-AP, 1:1000). Epicardial cells were maintained for up to 6 passages, as we noted decreased proliferation and altered cell morphology beyond this point.

### VCM differentiation

hiPSCs were differentiated to ventricular cardiomyocytes by temporal Wnt modulation followed by metabolic selection^21^. On day -3, hiPSCs were dissociated with accutase, and seeded at 75,000–150,000 cells per well of a Matrigel-coated 6-well plate (targeting ∼60–70% confluency at differentiation onset) in mTeSR+ with 10 µM Y-27632 (ROCK inhibitor). ROCK inhibitor was withdrawn the following day, and cells were maintained in mTeSR+ until day 0. On day 0, cells were switched to cardiomyocyte differentiation medium (CDM, see above) containing 6 µM CHIR99021 for exactly 48 h. On day 2, cells were rinsed with PBS and medium was replaced with CDM containing 5 µM IWP-2. Media was replaced with CDM alone on days 4 and 6. Spontaneous beating was reliably observed from day 6 onward.

On day 8, cardiomyocytes were dissociated with Accumax for 30-45 minutes, gently triturated, and replated 1:5 onto Matrigel-coated plates in cardiomyocyte maintenance medium (CMM, RPMI-1640 with 25 mM HEPES and GlutaMAX + B27 with insulin) with 20% knockout serum replacement (KOSR) and 10 µM Y-27632. Media was changed to fresh CMM without ROCK inhibitor on day 9. On day 11, cells were rinsed with PBS metabolically selected in cardiomyocyte selection medium (CSM; glucose-free RPMI-1640 with 5 mM sodium DL-lactate, 0.5 mg/mL BSA, and 213 µg/mL AA2P). CSM was refreshed every 48 hours for 8 days. Cells were recovered and maintained in CMM from day 19 onward. Differentiation efficiency was assessed by intracellular flow cytometry (see below) using anti-cardiac troponin-T (cTnT) antibody (BD 565744, 1:100).

### VCM differentiation competition assays

To assess relative differentiation efficiency of hiPSC genotypes, LMNA KO SLICE landing pad cells were mixed 1:1 with either LMNA CRISPR control WT cells or SLICE re-expressing WT LMNA cells on day -3 of VCM differentiation. Cells were differentiated in the same well for 12 days, then lifted with AccuMax and replated 1:5 for lactate selection and maturation to day 20. A portion of the Day 0, Day 12, and Day 20 cells were fixed and stained for LA/C (sc-376248 AF488, 1:250) and cTnT. The percentage of TnT+ cells was quantified for the LMNA- and LMNA+ cells in each well using FlowJo.

### Protein abundance screens

hiPSC library was expanded to 3 technical replicates of 10 million cells per N- and C-terminal antibody screen. Epicardial differentiation proceeded immediately after library generation in 3 technical replicates. For VCM differentiation, 2 frozen stocks of 2 million cells each (>200x library coverage) were thawed and recovered prior to seeding. The first biological replicate (3 technical replicates) was seeded one passage post-thaw, and the second replicate (2 technical replicates) was seeded one passage later. For all differentiations, 2 million library hiPSCs (>100x library coverage) were seeded per technical replicate. Cells expand during differentiation, so each replicate yielded 80 -100 million differentiated cells. Epicardial cells were collected on passages 3 and 4, while VCMs were collected on differentiation days 21 and 26. Cells were enzymatically dissociated with accutase (hiPSCs), TryPLE (epiCs), or accuMAX (VCMs), counted, and washed twice with TBS. The cells were fixed overnight at 4C in Z7 fixative (0.5% zinc chloride, 0.5% zinc trifluoroacetate, 0.05% calcium acetate in 0.1M Tris pH 6.8), then washed twice with TBS. Cells were permeabilized and blocked in staining buffer (1% FBS, 0.1% TritonX-100, 0.02% SDS in TBS) for 30 minutes at room temp, then co-stained with conjugated LMNA/C N-terminal antibody (E-1, Santa Cruz Biotechnology, sc-376248 AF488, 1:250) and lineage marker antibodies (WT-1 for epicardial cells, cTnT for VCMs) for 1 hour at room temp, rocking. For the C-terminal LMNA hiPSC screen, cells were incubated at 4C overnight with primary antibody (Sigma L1293, 1:500), washed twice with staining buffer, and then incubated for 1 hour with AF488-conjugated secondary antibody. Cells were washed twice in staining buffer, then once in TBS, and resuspended to 10 million cells per mL for FACS. Cells were strained through 40 µm (hiPSC, EpiC) or 70 µm (VCM) strainers prior to sorting on a BD Fusion cell sorter. The cells were first gated on their respective lineage marker, and then the lamin intensity histogram was gated into 4 bins, each containing 25% of the total signal. For each replicate, 500,000 to 1 million cells were collected per bin into 5 mL tubes containing 750 uL concentrated FACS buffer (10% FBS, 5 mM EDTA in TBS). Cells were transferred to 15-mL conical tubes, pelleted at 1200 g for 5 minutes, then resuspended in 100 uL TBS. Amplicon sequencing was performed as described above, and variant counts were analyzed using Lilace^26^ (v1.0.0) with positional shrinkage enabled.

### VCM fitness screen

To obtain library representation counts between epiC and VCM pools, aliquots of 2 million unsorted cells were taken from each unsorted EpiC and VCM differentiation technical replicate. 5 samples per cell type were processed alongside sorted cell pools (see above).

The counts files were fed into Rosace^49^ (v1.1.0), with the epicardial replicates set as AssaySet time = 0 and the VCM replicates set as time = 1. Consistent with the variable nature of hiPSC differentiation, the Rosace output was noisy: 92% of the variants had a 95% CI overlapping the synonymous null distribution, and the median per-variant uncertainty (SD 0.56) was nearly as large as the spread of scores across the entire dataset (SD 0.84; ∼67%). Therefore, only variants that met all of the following criteria were classified as “depleted”:

Called significanly negative by Rosace: Label = “Neg”, Rosace LFSR (Local False Sign Rate) < 0.05
Upper bound of the 95% confidence interval (Rosace score + 1.96 x SD) fell below the 5^th^ percentile of the synonymous score distribution
Located at positions where the mean Rosace score was <1.5x the library mean
Located at positions where the fraction of significantly depleted (Rosace LFSR < 0.05) >1.5x the library mean

We classified these variants as “expressed” if they were 1) not called significantly depleted by Lilace, and 2) their abundance scores were greater than the 5^th^ percentile of synonymous abundance scores. For comparison to human population databases, we loosened the positional filters to 1x, which yielded a list of 533 low-fitness variants.

### Immunoblotting for Lamin A expression

Cell pellets were resuspended in Urea Lysis Buffer (8 M urea, 75 mM NaCl, 50 mM Tris pH 8.0 and cOmplete, Mini, EDTA-free Protease Inhibitor Cocktail (Roche, 11836170001). Lysates were sonicated with 2x 10-second pulses on a probe sonicator set to 30% power, then spun at max speed for 2 minutes to pellet any remaining insoluble material. An aliquot of lysate was diluted 1:5 and quantified by BCA (Pierce). 20-50 ug protein was resolved on a 10% SDS-PAGE gel and transferred to a nitrocellulose membrane. Blots were probed with antibodies against the Lamin A/C Ig-Fold (Active Motif 3A6-4C11, 1:1000) or N-terminus (Santa Cruz Biotechnology sc-376248, 1:1000), alongside GAPDH (GeneTex gtx300041, 1:5000) or tubulin (Sigma-Aldrich T5168, 1:2000) as loading controls. Blots were then washed and incubated with fluorescently conjugated secondary antibodies prior to imaging on a Licor Odyssey instrument.

### Nuclear fractionation by high-salt extraction

Lamin A/C variants were expressed in *LMNA*-knockout hiPSCs via the SLICE landing-pad system. For each sample, 2 × 10⁶ cells were pelleted, snap-frozen, and lysed by freeze/thaw. Pellets were resuspended in isotonic buffer (10 mM HEPES pH 7.4, 2 mM MgCl₂, 25 mM KCl, 250 mM sucrose, protease inhibitors); an aliquot was retained as whole-cell lysate, and nuclei were pelleted (376×g, 5 min, 4 °C) to yield the cytosolic fraction. Nuclei were then extracted in high-salt buffer (20 mM HEPES pH 7.4, 1 M NaCl, protease inhibitors) for 15 min at room temperature with agitation, and centrifuged (10,000×g, 5 min, 4 °C) to separate the soluble fraction from the insoluble pellet, which was solubilized directly in SDS-PAGE sample buffer. Equal volumes of each fraction were resolved by SDS-PAGE and immunoblotted for Lamin A/C, GAPDH (cytosolic marker), and histone H3 (chromatin marker). Variants released into the soluble fraction with NaCl were interpreted as less stably incorporated into the insoluble lamin meshwork.

### Immunofluorescence microscopy

Cardiomyocytes were replated for imaging as follows: cells were dissociated with Accumax for 30-45 minutes, counted, and re-seeded at 40,000 cells per well of a Matrigel-coated ibidi micro-Slide chamber (ibidi, 80806) in CMM + 20% KOSR and ROCK inhibitor. Media was replaced with fresh CMM the next day, and cells were allowed to recover for 4 days before fixation.

Epicardial cells were seeded on gelatin-coated ibidi dishes in LaSR replating medium at 10,000 cells per well, and media was replaced with fresh LaSR the next day. Cells were allowed to expand in the ibidi well for an additional 2 days before fixing.

hiPSCs were seeded on Matrigel-coated ibidi dishes at 50,000 cells per well in mTeSR+ with ROCK inhibitor the day prior to fixation. Cells were left in ROCK inhibitor until fixation to promote monolayer adhesion.

Cells were fixed in 4% PFA/PBS for 10 minutes at room temperature, washed twice in PBS, and permeabilized/blocked in IF buffer (1% FBS, 0.1% TritonX-100, 0.02% SDS in PBS) for 30 minutes at room temperature. Primary antibodies were diluted in IF buffer and added to cells for 1-2 hours at room temp or 4C overnight. Cells were washed 3x with IF buffer and incubated with conjugated secondary antibodies and Hoechst (10 ug/mL) in IF buffer for 30 minutes to 1 hour, room temperature. Cells were then washed 2x with IF buffer and once with PBS. After washing, PBS was added to ibidi wells for imaging.

### Image acquisition and processing

Confocal z-stacks (step size 0.3 µm) were acquired on a Nikon spinning disk confocal microscope with a 40x/0.95 NA or 60x/1.4 NA objective. 189-363 cells were imaged per genotype across multiple independent imaging experiments. Unprocessed 16-bit images were saved as ND2 files using the Nikon Elements 5.02 build 1266 software. Pixel size and z-step were read from the file metadata.

Images were analyzed using a custom Python pipeline. Nuclei were segmented from the Hoechst (DNA) channel. The 3D stack was blurred to reduce noise (Gaussian filter, σ = 1.5 pixels in xy and 1.0 in z), Otsu-thresholded, and the resulting mask cleaned by 3D morphological closing then opening (spherical element, radius 2 voxels) to fill small holes and remove specks. To split touching nuclei, the mask was collapsed to a maximum-intensity projection and each pixel labeled by its distance to the nearest edge (Euclidean distance transform, smoothed with σ = 3); the peak of this distance map within each nucleus defined its center. These centers seeded a marker-controlled watershed, which places boundaries between adjacent nuclei. Seeds were required to be ≥ 6 µm apart — specified in physical units and converted to pixels for each image, so the spacing is consistent across magnifications and elongated nuclei are not over-split. The 2D outlines were then propagated through the z-planes to reconstruct each nucleus in 3D. Border-touching nuclei were excluded, as were objects outside 50–500 µm² (30–800 µm² for pixel sizes < 0.2 µm). Under-segmented or contacting nuclei were removed by a boundary check that excluded any nucleus whose mean Hoechst intensity in a 3-px perinuclear ring exceeded 0.5× (0.9× at high magnification) its interior mean.

To assess nuclear morphology we measured cross-sectional area, perimeter, circularity, deformation (1 − circularity), solidity, eccentricity, aspect ratio, and 3D volume/height. Contour regularity was quantified as an elliptic Fourier coefficient (EFC) ratio (15 harmonics; first-harmonic semi-axes ÷ summed higher-harmonic semi-axes; higher = more regular).

To quantify Lamin A radial distribution, the central (widest) z-slice of each nucleus was chosen; its mask was solidified and dilated by 1 px to capture the lamina just beyond the chromatin edge. A Euclidean distance transform was normalized from 0 (center) to 1 (edge) and divided into 25 concentric shells; each shell’s Lamin A signal was expressed as fractional intensity (shell mean ÷ whole-nucleus mean). To exclude mitotic/mis-segmented nuclei from radial intensity analysis, nuclei with an EFC ratio < 2 and cells with the lowest 20% nuclear area were discarded prior to calculation.

On the same central z-slice, an emerin nuclear-rim fraction was computed as the fraction of local emerin signal within a 5-px-wide perinuclear rim (the nucleus minus its 5-px erosion) relative to a local cell region (the nucleus dilated by 40 px, excluding neighboring nuclei); the ratio of rim to cytoplasmic mean intensity was also recorded.

### Data analysis

Lamin A protein sequence was obtained from Uniprot ID P02545, human Prelamin-A/C and used for library design and all subsequent analysis. Lamin A domain boundaries were assigned according to UniProt annotation: head 1–33; coil 1A 34–70; linker 1 71–80; coil 1B 81–218; linker 2 219–242; coil 2 243–383; tail 384–416; NLS 417–420; tail-2 421–429; Ig-fold 430–546; tail-3 547–664. Mutant residues were classified as follows: positive: R, H, K; negative: D, E; polar: S, T, N, Q; special: P, G, C; hydrophobic: A, V, I, L, M; aromatic: Y, W, F; indel: insertion G/GS/GSG, deletion 1, 2, 3.

Analyses were performed in GraphPad prism (v10.6.1) and R (v4.3.3) using rstatix v0.7.2 and binom v1.1-1.1 (statistical testing and confidence intervals), and ggplot2 v4.0.3 for visualization.

### Evolutionary analysis

Per-residue evolutionary conservation of Lamin A/C was computed with the ConSurf web server^64^ (https://consurf.tau.ac.il/) using the human Lamin A/C sequence (UniProt P02545) as the query. Homologs were collected with the default search (one HMMER iteration against UniRef90, E-value cutoff 1×10⁻⁴, up to 150 sequences). The MSA was built with MAFFT, and position-specific conservation scores were plotted directly or normalized to a nine-grade scale (1, most variable; 9, most conserved). Because no structure was supplied, buried/exposed and functional/structural residue assignments were derived from the AlphaFold model automatically retrieved for P02545.

To assess residue coevolution, human Prelamin-A/C (Uniprot ID P02545) was used as bait sequence for EVcouplings analysis^48^ with default settings (5 iterations, pseudo-likelihood maximization inference method). A multiple sequence alignment (MSA) was built by jackhammer against UniRef90, and was limited to the lamin rod domain (positions 30 to 386). Sequences with a bitscore threshold of 0.3 bits/residue were selected, yielding 4523 aligned sequences (14.4 effective sequences per residue; EVcouplings alignment quality 5/10).

### Structural modeling and thermostability predictions

Coiled-coil propensity and heptad register of Lamin A were predicted with MARCOIL^65^ via the MPI Bioinformatics Toolkit server (https://toolkit.tuebingen.mpg.de/tools/marcoil), using the full-length Lamin A sequence (UniProt P02545) with default settings. Per-residue coiled-coil probability and heptad register (a–g) were retained where register probability exceeded 80%, with probabilities scaled to 0–1 for visualization. Register assignments were used to map heptad phase along the rod domain and locate coiled-coil discontinuities, including the coil-2 stutter (positions 327–337) and a putative linker within coil 2 (positions 278–288). Positions were manually annotated for hendecad repeats according to^44^.

The ACN tetramer model was generated using the AlphaFold3 server^47^. Two N-terminal chains (A,B) corresponding to Lamin A residues 1-70 and two C-terminal chains (C,D) corresponding to residues 327-401 were submitted. The first of 4 output models (model_0) was selected for further analyses. UCSF ChimeraX-1.10 was used for structure visualization. DMS scores were mapped onto ACN and A11 (PDB 6JLB) structures using the mutationscores Chimera command.

For analysis of whole protein stability, a model of the Lamin A dimer was generated using AlphaFold2 ColabFold and fed into the Colab ThermoMPNN implementation^66^ (ThermoMPNN.ipynb) with cysteines excluded and multichain inference enabled.

The energetic consequences of LMNA variants to dimer/tetramer structures (A11, PDB 6JLB; ACN, AlphaFold3 model) were estimated using FoldX5^33^. Starting from each homodimer model, side-chain geometry was first optimized with the RepairPDB command. For each mutation, mutant structures were generated with the BuildModel command using three independent runs, and the resulting complexes were evaluated with the AnalyseComplex command across the two interchain interfaces (chains AB and CD) to obtain interaction energies. Per-mutation BuildModel and AnalyseComplex outputs were collated into position-specific scoring matrices (PSSM) of ΔΔG values (in kcal/mol) for every amino acid substitution. Calculations were performed on both the dimer and tetramer (anti-parallel/ACN and A11) interface models so that the differential effect of each variant on each interface could be assessed. A variant was classified as destabilizing for a given interface when its ΔΔG exceeded 1.6 kcal/mol, and each variant was assigned to one of four categories—“dimer,” “tetramer,” “both,” or “neither”—based on whether it destabilized the dimer interface, the tetramer interface, both, or neither.

Random forest regression models were trained to predict the per-variant VCM abundance score for Lamin A missense and synonymous variants to assess how well sequence- and structure-derived features predict abundance scores. Predictors comprised WT and mutant amino acid identity, mutation type, WT and mutant amino acid biochemical class, biochemical class transition, domain, residue position, per-residue evolutionary conservation score from ConSurf^64^, buried/exposed status (ConSurf predicted), BLOSUM62 substitution score^67^, and predicted folding stability change (ΔΔG) from ThermoMPNN^66^. Positions 1–7 were excluded (antibody epitope region). Random forests were grown with the ranger package^68^ (v0.17.0) in R using 500 trees, default settings, and ordered handling of unordered categorical predictors. Model performance was assessed on held-out data as the out-of-bag R² and the 5-fold cross-validated R² (reported as mean ± SD across folds). Feature importance was estimated by permutation.

### Clinical variant comparisons

Human *LMNA* variant classifications were downloaded from ClinVar (accessed 21 December 2025; *LMNA* gene query). Protein-level changes were parsed from the variant Name field and converted to single-letter amino-acid codes; entries lacking a valid protein-level consequence were discarded. Germline classifications were collapsed into P/LP (“Pathogenic,” “Likely pathogenic,” “Pathogenic/Likely pathogenic”), B/LB (“Benign,” “Likely benign”), VUS (“Uncertain significance,” “Uncertain significance/Uncertain risk allele”), conflicting (“Conflicting classifications of pathogenicity”), and not provided (missing or “not provided”). Variants were not filtered by review status (star rating). ClinVar variants were matched to the DMS library by protein-level change to associate each with its abundance and fitness scores.

### Human population database analysis

To test whether functionally important variants were under-represented in human population databases, we compared variant sets (variants depleted in the VCM fitness screen or lowly expressed in the abundance screen) against all other non-indel library variants by presence in the combined gnomAD^35^ (accessed 7/21/25) and Regeneron datasets^36^ (accessed 7/21/25), using one-sided Fisher’s exact tests (alternative = “less”) for depletion. Indels are sparsely represented in population databases and were thus excluded; variant set proportions are reported with 95% Wilson confidence intervals.

Because mutation classes (missense, nonsense, synonymous) occur at systematically different population frequencies, we controlled for composition by permutation (1,000 iterations): in each, a random variant set matching the focal set’s mutation-type composition was drawn from the library and subjected to the same test. Depletion was called significant when the observed focal-set proportion fell below the 2.5th percentile of the permutation null.

Cardiac phenotype penetrance was assessed from the All of Us research study (allofus.nih.gov^37^). Genomic and electronic medical record data from 535,662 study participants in the Curated Data Repository version 9 were queried for LMNA CDS mutations and cardiac-related ICD-10 and ICD-9 codes (corresponding to atrioventricular (AV) block, arrhythmogenic cardiomyopathy (ACM), atrial fibrillation (AFib), cardiac arrest, cardiac transplant, dilated cardiomyopathy (DCM), heart failure, implantable cardiac defibrillator (ICD) or pacemaker, ventricular arrhythmia). Samples that were QC-flagged, had genotype quality <20, or read depth <10 (1,012 participants total) were excluded. 2,977 individuals were identified carrying 359 unique nonsense or missense mutations in the Lamin A structured domains (residue positions 8-547). These variants were matched with scores from 13,265 missense, nonsense, and synonymous variants measured in the VCM protein abundance screen. Variants were called low abundance if their score was less than the 5^th^ percentile of the synonymous distribution. Penetrance was calculated by dividing the number of individuals with a given ICD-10 or ICD-9 diagnosis by the number of genotype positive individuals in the same stratification (low abundance variants, other LMNA variants that were not low abundance, and the total All of Us population). Analyses were run in the All of Us Researcher Workbench.

### OddsPath calculation for functional evidence strength

To assess whether LILACE abundance scores provide evidence of pathogenicity suitable for clinical variant interpretation, we computed OddsPath ratios following the ClinGen Sequence Variant Interpretation Working Group framework^38^. Pathogenic controls were ClinVar P/LP missense variants (n = 195). Benign controls were Clinvar B/LB variants and missense variants observed in gnomAD and/or the Regeneron exome dataset at allele frequencies >2×10⁻⁵ that were not classified as P/LP in ClinVar and were present in our DMS library (*n* = 43, supp. table). P1 was defined as the proportion of pathogenic variants among all controls, and P2 as the proportion of pathogenic variants among those scoring below the 12.5^th^ percentile of the synonymous score distribution, with Laplace smoothing (P2 = (n pathogenic below + 1) / (n pathogenic below + n benign below + 2)). OddsPath was calculated as [P2 × (1 − P1)] / [(1 − P2) × P1]. Evidence strength was assigned per established thresholds: OddsPath >2.1, PS3_supporting; >4.3, PS3_moderate; >18.7, PS3_strong. OddsPath was evaluated across a range of score thresholds to identify the optimal threshold and across allele frequency thresholds for benign control definition to assess sensitivity to this parameter.

## Author Contributions

J.M., W.C-M., and A.B. conceptualized the study; J.M., A.H., N.L., and J.C. performed experiments; J.M., A.H., J.C., and C.Y. analyzed data; J.M. and A.B. wrote the manuscript; and A.L., V.V., and A.B. supervised project progress and provided funding support.

---

**Figure 1 supplement 1.**
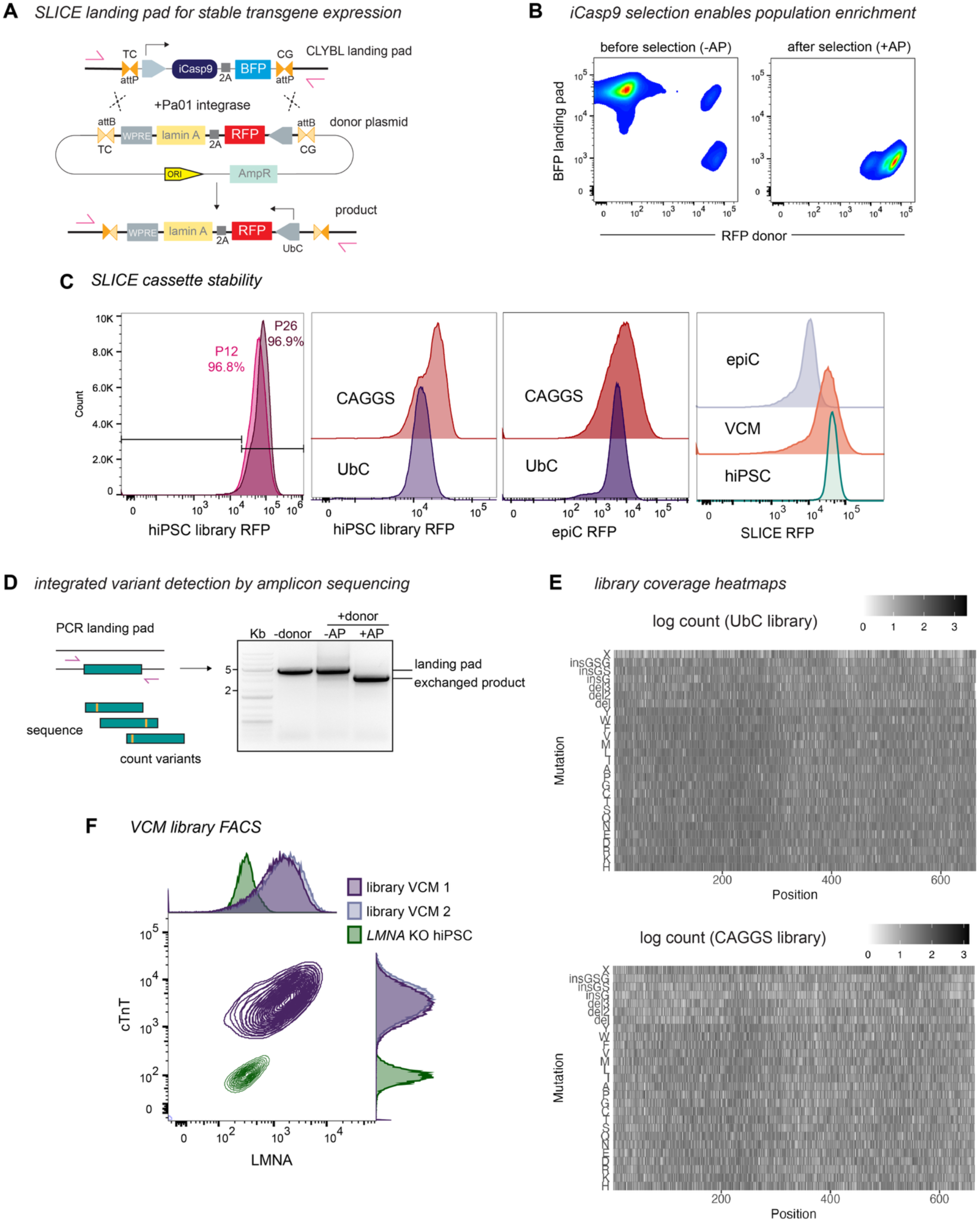
SLICE platform for DMS library integration. A) SLICE diagram. Landing pad with BFP and iCasp9 negative selection marker flanked by two heterotypic Pa01 recognition sites (attP, TC or CG central dinucleotide pair) in CLYBL safe harbor locus. Transfection with attB-containing donor plasmid and Pa01 results in directional cassette exchange. Arrows, genotyping primers. B) Example selection FACS plots of landing pad cells transfected with RFP-containing donor plasmid before (left) and after (right) iCasp9 activation with AP1903 (AP). C) FACS plots showing SLICE cassette expression over time, between promoters, and across cell states. SLICE cassette mCherry (upstream of P2A-lamin A) fluorescence quantified across conditions. Left: FACS plot of hiPSC CAGGS library, 12 and 26 passages after SLICE integration. Center left: CAGGS versus UbC library expression in hiPSCs. Center right: CAGGS versus UbC library expression in epiCs. Right, fluorescence quantified from isogenic hiPSCs, epiCs, and VCMs. SLICE cassette expression is slightly decreased after epicardial differentiation. D) To quantify variants from the SLICE landing pad, genomic DNA from landing pad cells ± donor plasmid, ± AP1903 selection was amplified with primers shown in (A). Selection results in complete loss of upper landing pad band. Exchanged product is excised, purified, and sequenced to count integrated variants. E) Heatmaps showing log10 counts of lamin A variant libraries (UbC promoter, top; CAGGS promoter, bottom) after SLICE integration. x-axis, lamin A sequence position; y-axis, variant mutations. F) VCM library lamin A protein abundance and cardiac troponin T (cTnT) expression. Lamin A variant library VCMs and *LMNA* KO hiPSC controls were stained for LMNA/C and cTnT. VCM library was uniformly cTnT+, while LMNA/C abundance varied across the population. For abundance screen, cells were gated on cTnT+ and sorted on LMNA/C fluorescence intensity.

**Figure 1 supplement 2.**
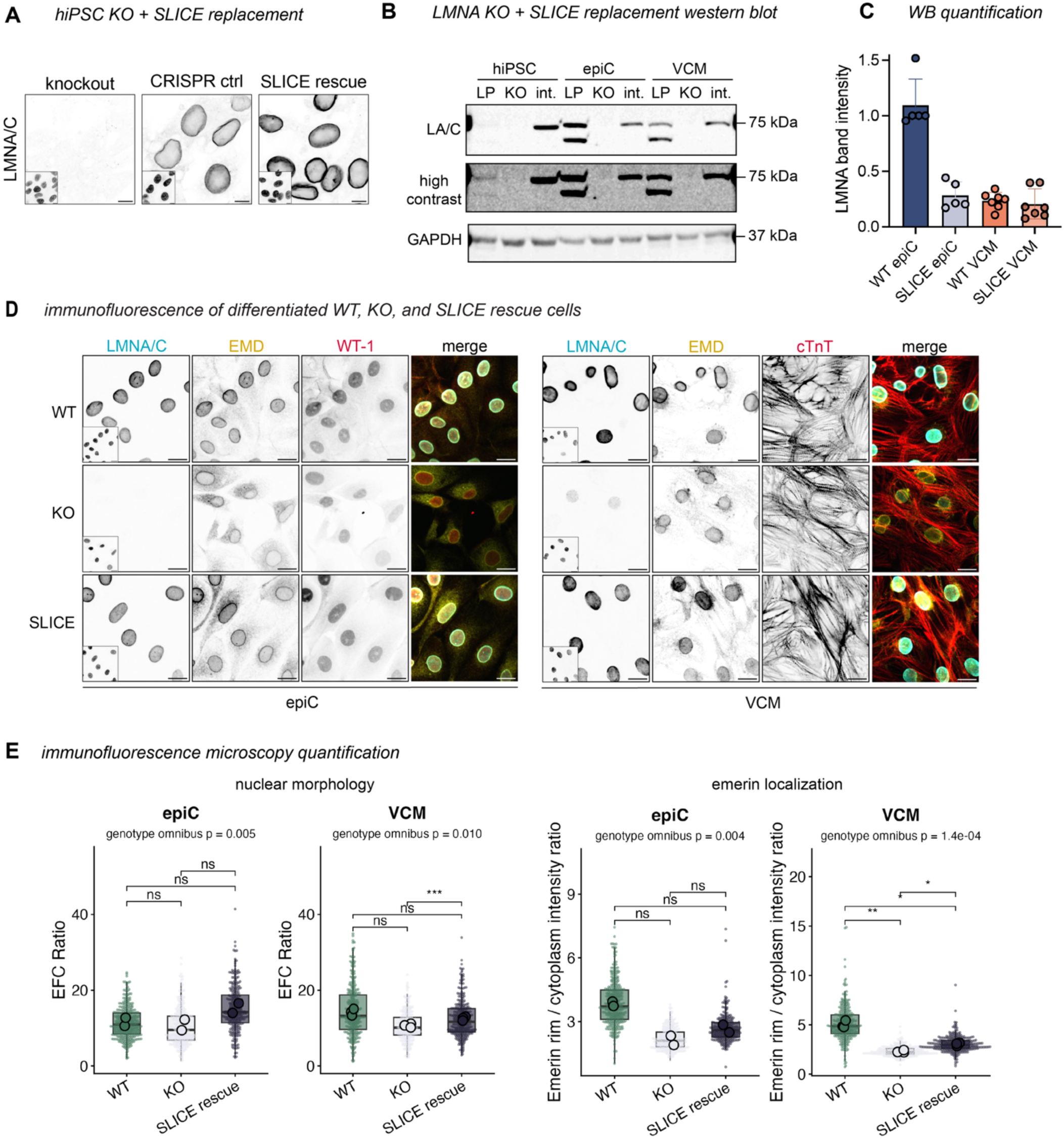
Lamin A expression from SLICE landing pad. A) Immunofluorescence of *LMNA* KO, CRISPR control WT, and *LMNA* KO SLICE landing pad + re-integrated WT lamin A hiPSCs, stained for LMNA/C. Inset, Hoechst DNA stain. Scale bar, 10 µm. B) Example Western blot of LMNA/C levels in *LMNA* WT SLICE landing pad (LP), *LMNA* KO, and *LMNA* KO SLICE landing pad + re-integrated WT lamin A (int.) hiPSCs, epiCs, and VCMs. High contrast panel shows low level expression of endogenous LMNA/C in hiPSCs. GAPDH, loading control. C) Quantification of Western blots exemplified in (B). Band intensity was normalized to one epiC WT lamin A band (top) per blot. *n* = 7 VCM, 5 epiC samples across 4 blots. D) Immunofluorescence images of WT landing pad cells, *LMNA* KO landing pad cells, and *LMNA* KO landing pad cells SLICE-integrated with WT *LMNA*, differentiated into epicardial cells (epiC) or ventricular cardiomyocytes (VCM). Stained for LMNA/C, emerin (EMD), and lineage marker (WT1, epiC; cTnT, VCM). Scale bars, 20 µm. E) Quantification of imaging experiments. Box plots show nuclear area (left), EFC ratio (center, higher = more regular nuclear morphology), emerin nucleus:cytoplasm intensity ratio (right). Small points, individual nuclei (box = pooled median and IQR); large points, per-replicate means (unit of *n*). Omnibus *P*: nested ANOVA (*n* = 2 epiC / 3 VCM biological replicates) indicates difference between all cell types. Brackets: Holm-corrected pairwise paired *t*-tests on replicate means (\**P* < 0.05, \*\**P* < 0.01, \*\*\**P* < 0.001).

**Figure 1 supplement 3.**
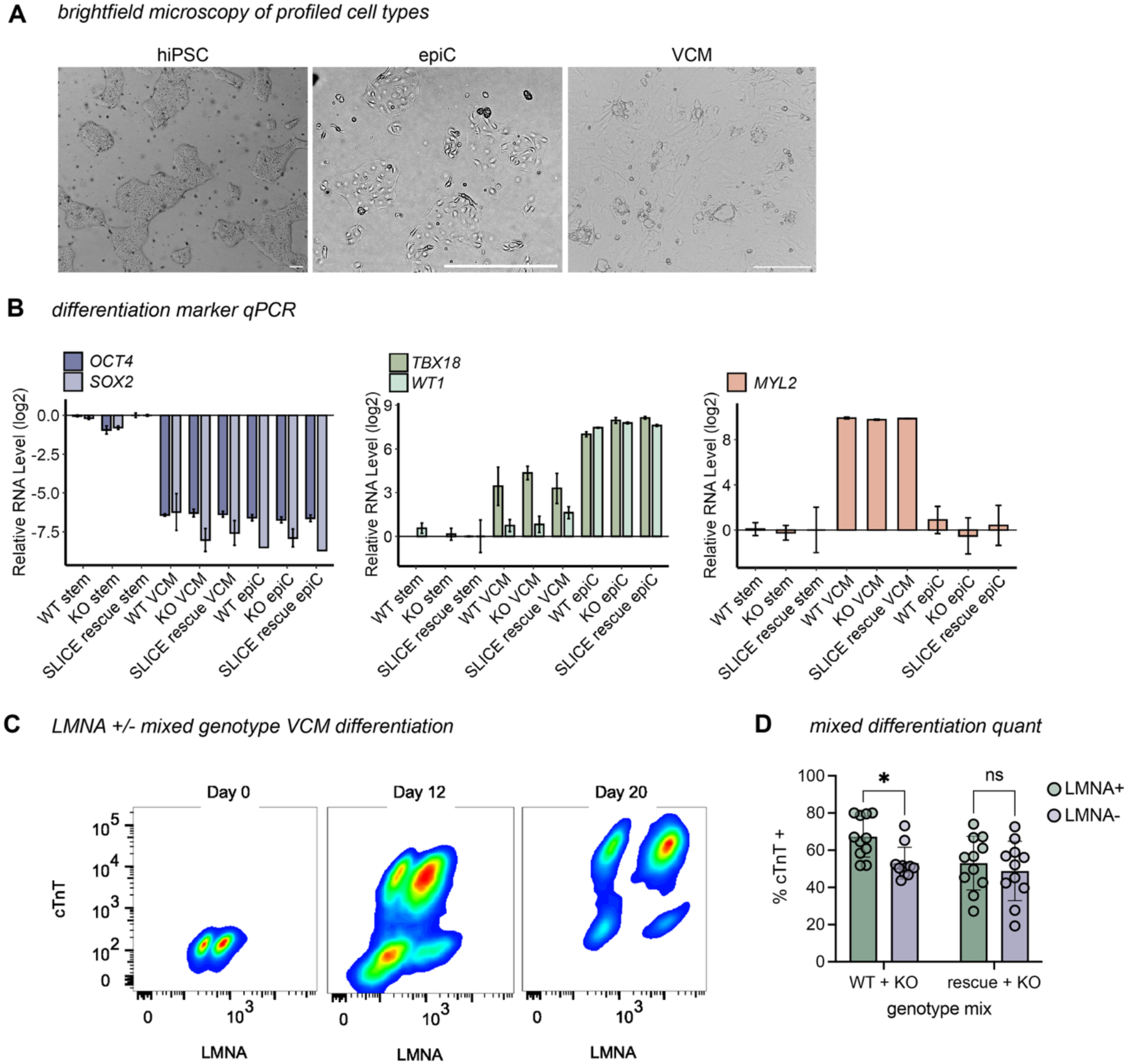
Differentiation of SLICE landing pad hiPSCs. A) Brightfield microscopy of hiPSCs, epiCs, and VCMs showing characteristic morphology of each cell type. Scale bars, 200 µm. B) qPCR for lineage-specific genes in *LMNA* WT SLICE landing pad (WT), *LMNA* KO, and *LMNA* KO SLICE landing pad + re-integrated WT lamin A (SLICE rescue) hiPSCs, epiCs, and VCMs. *OCT4* and *SOX2*: hiPSC markers, *TBX18* and *WT1*: epicardial markers, *MYL2*: VCM marker. Transcript levels are normalized to WT hiPSC condition and log2 scaled. C) FACS plots of LMNA/C and cardiac troponin T (cTnT) expression over VCM differentiation. WT and *LMNA* KO cells were mixed 1:1 and seeded in the same dish at differentiation day −3. Cell aliquots were collected at days 0, 12, and 20, fixed, permeabilized, and stained for LMNA/C and cTnT to assess differentiation efficiency. LMNA/C+ cells reliably produced higher levels of cTnT, while cTnT− cells were more likely to be *LMNA* KO. D) Quantification of mixed differentiation experiments exemplified by (C). *LMNA* KO cells were either mixed with WT SLICE landing pad hiPSCs (WT + KO, left bars) or *LMNA* KO with WT lamin A integrated into the SLICE landing pad (rescue + KO, right bars). At day 12, the percentage of LMNA/C+ and LMNA/C− cells that expressed cTnT was quantified by flow cytometry. SLICE re-expression of lamin A does not rescue KO differentiation defect. \**P* = 0.003 by unpaired *t*-test with Bonferroni adjustment. *n* = 11 separate mixed wells over 4 independent experiments.

**Figure 1 supplement 4.**
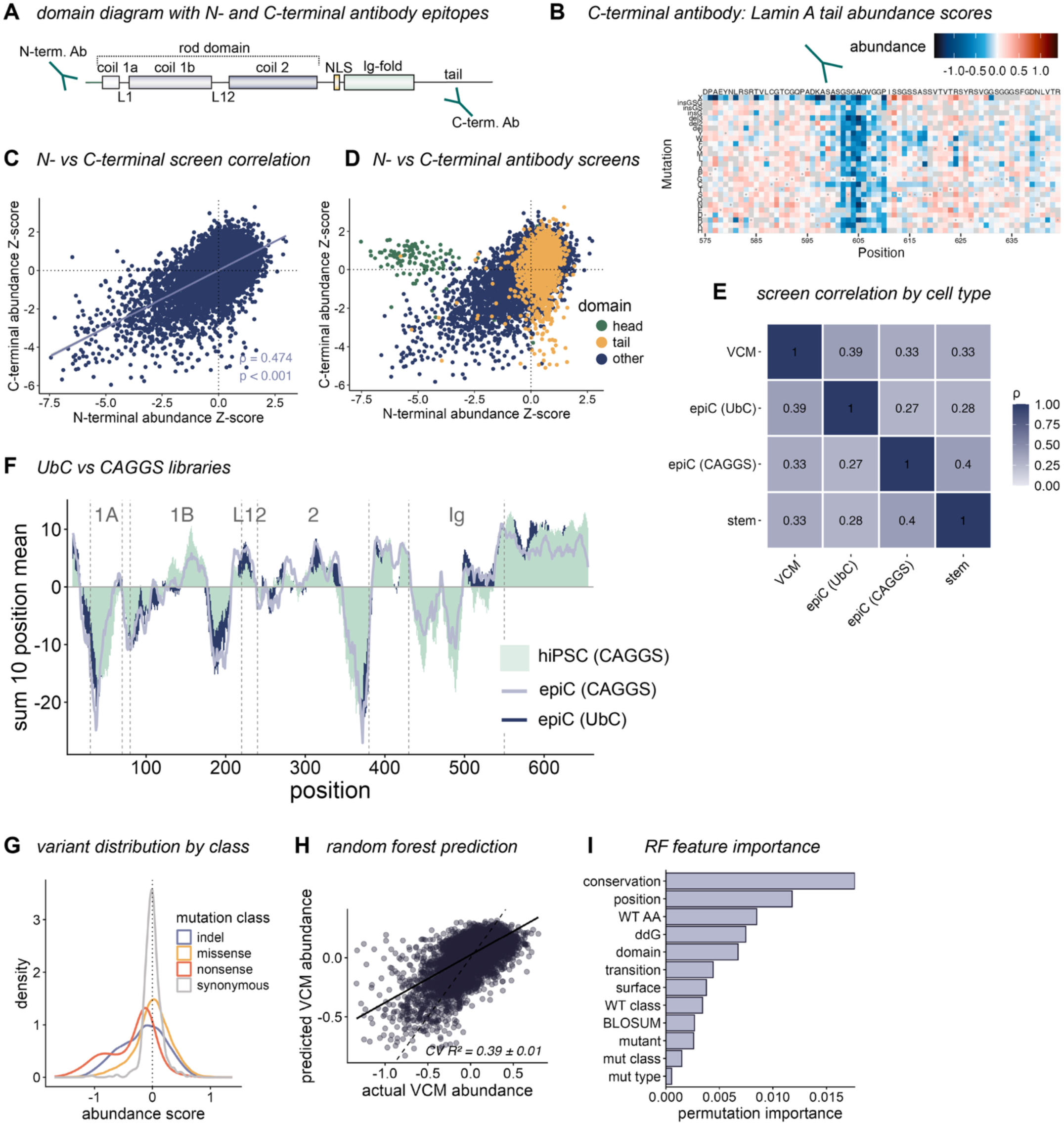
Comparisons of immunostaining screens. A) Diagram of lamin A domains and antibodies used for protein abundance screens. B) Heatmap segment (residues 575–664, tail domain) of abundance scores from C-terminal antibody screen in hiPSCs. Prominent vertical stripe in C-terminal heatmap not present in N-terminal heatmap, thus mapping C-terminal epitope to residues 602–610. C) Correlation between z-transformed abundance scores for hiPSC screens with N- and C-terminal antibodies, with epitopes excluded. Blue line, best fit. Spearman’s rho indicated at bottom left. D) Comparison of z-transformed abundance scores for hiPSC screens with N- and C-terminal antibodies, colored by domain. A subset of head and tail domain residues only display low signal in N- and C-terminal screens, respectively. E) Spearman correlation heatmap for N-terminal antibody abundance screens in hiPSCs (stem, CAGGS promoter), epiCs (two promoters), and VCMs (UbC promoter). F) Rolling-window abundance signal along lamin A by promoter and cell type. Each trace is the rolling sum of z-scored position-mean Lilace abundance scores over 10-residue windows (1-position step), plotted at the window center, for each library. Relative effect differences between hiPSC library and epiC libraries are independent of transgene promoter. G) Example density plot of DMS screen data. Distribution of variant abundance differs by mutation class, with synonymous scores centered at zero and nonsense scores skewing negative. H) Random forest model predicted versus observed VCM abundance scores for the lamin A protein, with each point indicating one variant. Dashed line, identity (y = x); solid line, linear best fit with its 95% confidence interval (grey band). The model explains 39% of the variance in abundance (5-fold cross-validated R²). I) Permutation feature importance for the random forest (RF) model in (H), ranked by the decrease in predictive accuracy when each feature is shuffled. *n* = 10,378 variants.

**Figure 1 supplement 5.**
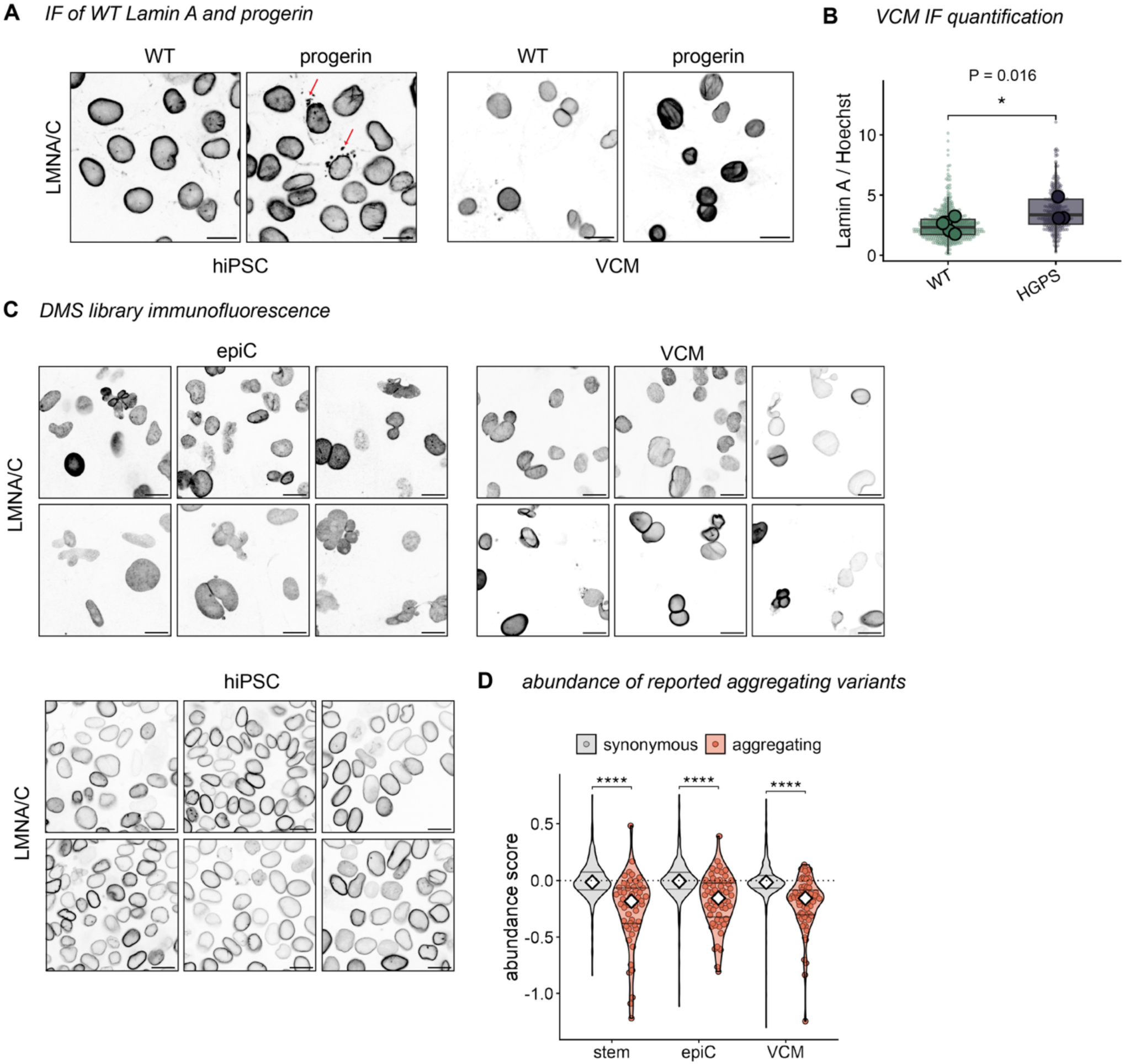
Protein aggregation is not a common outcome of lamin A mutation. A) anti-LMNA/C immunofluorescence micrographs of *LMNA* KO hiPSCs (left) and VCMs (right) expressing WT lamin A or progerin from the SLICE landing pad. Red arrows indicate patterns of cytoplasmic aggregates not noted in the WT cells. Scale bars, 20 µm. B) Quantification of LMNA/C fluorescence intensity normalized to Hoechst DNA stain for VCMs expressing WT lamin A or progerin from the SLICE landing pad. Significance calculated by paired *t*-test on the means of 3 independent imaging experiments. C) anti-LMNA/C immunofluorescence micrographs of lamin A DMS library in hiPSCs, epiCs, and VCMs. Six representative fields of view per cell type. Nuclear morphology defects are more apparent in differentiated libraries, while aggregate foci are rarely noted. Scale bars, 20 µm. D) Abundance scores across cell types for synonymous variants (grey; *n* = 636) and missense variants previously reported to significantly aggregate when GFP-tagged and overexpressed in C2C12 mouse myoblasts (orange; *n* = 66, Anderson et al., ref. 31). Violins show the distribution; white diamonds mark medians; points are individual aggregating variants. \*\*\*\**P* < 0.0001 by Wilcoxon rank-sum tests with FDR adjustment.

**Figure 3 supplement 1.**
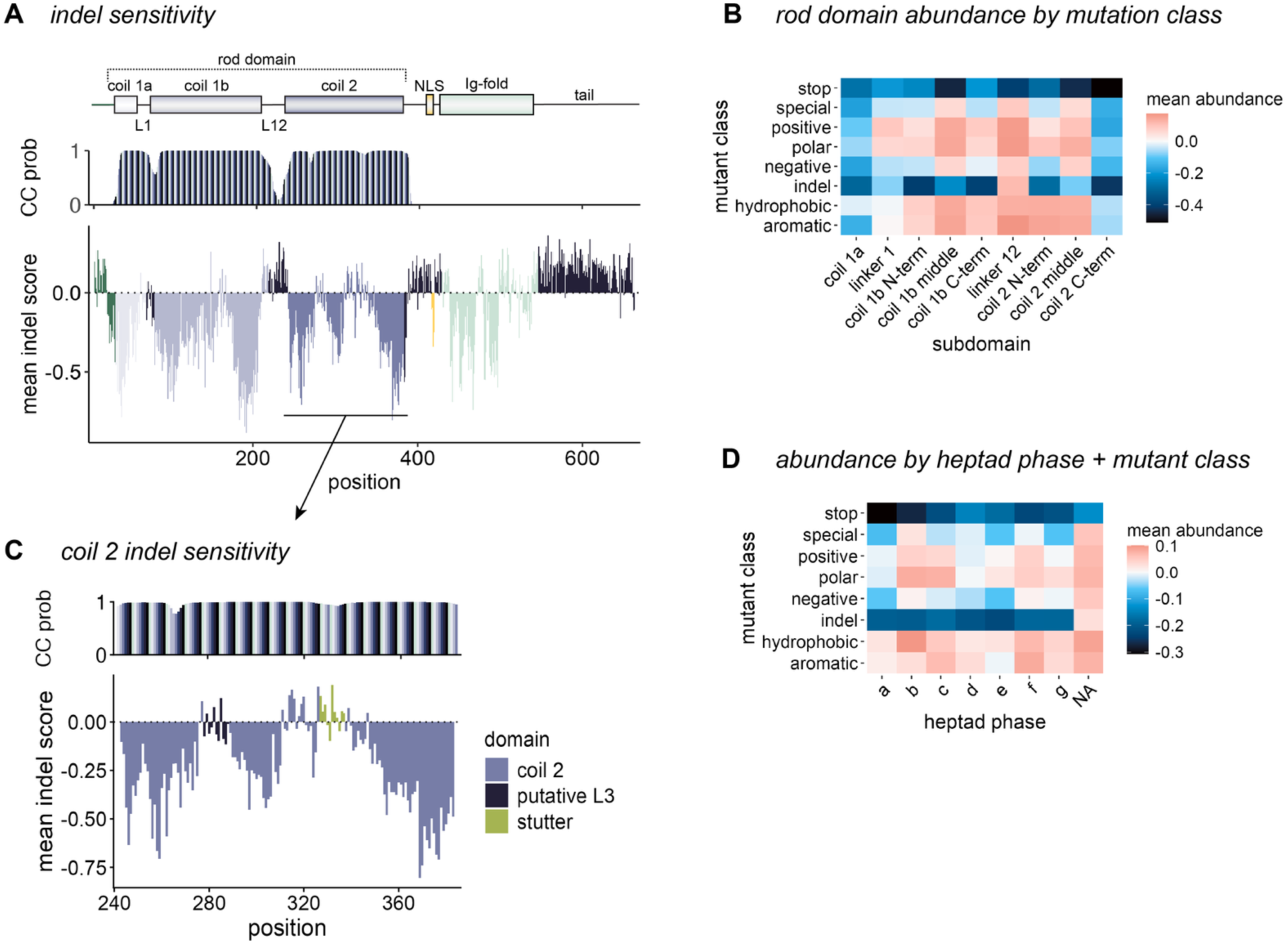
Mutation sensitivity of coiled-coil rod domain. A) Top: lamin A domain diagram with rod subdomains (coils 1a, 1b, coil 2, linkers L1 and L12) annotated. NLS, nuclear localization signal. Middle: CC prob, coiled-coil probability calculated with MARCOIL; heptad repeats indicated by stripes in probability plot. Bottom: mean VCM abundance score for all indels at each position in lamin A sequence, colored by domain. B) Heatmap of mean abundance scores for each mutation class (see Methods) across rod subdomains (coil 1a: positions 28–70, linker 1: 71–80, coil 1b N-term: 81–115, coil 1b middle: 116–170, coil 1b C-term: 171–218, linker 12: 219–242, coil 2 N-term: 243–280, coil 2 middle: 281–349, coil 2 C-term: 350–388). C) Zoomed mean VCM abundance score for all indels at each position within coil 2. Putative linker L3 (positions 278–288) and stutter sequence (positions 327–337) highlighted by color. D) Heatmap of mean abundance scores for each mutation class by heptad phase. NA, all positions outside rod domain.

**Figure 3 supplement 2.**
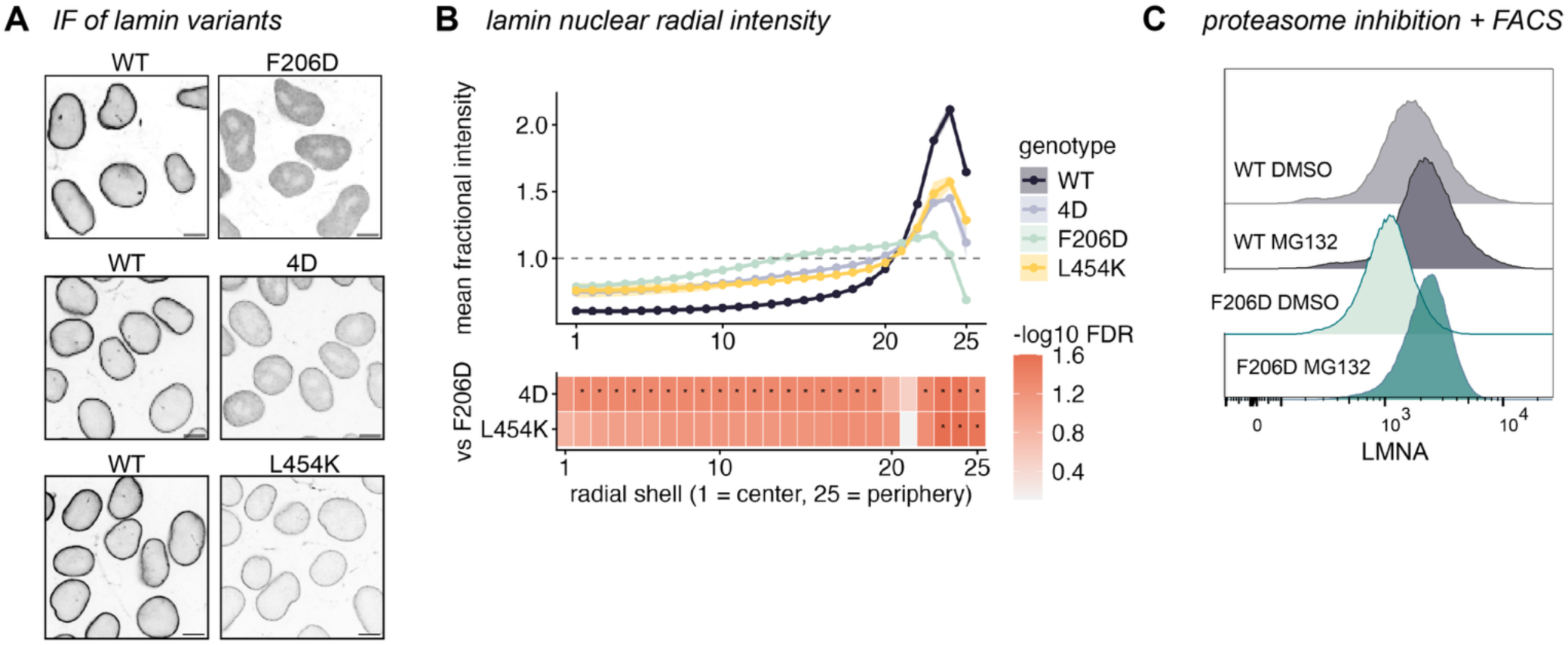
Mutation sensitivity of A11 multimer interface across cell types. A) Immunofluorescence images of *LMNA* KO cells expressing indicated mutants in SLICE landing pad, stained with N-terminal LMNA/C antibody. Scale bar, 10 µm. B) Fractional intensity (shell mean ÷ whole-nucleus mean) from nuclear center (shell 1) to periphery (shell 25). Lines are means across biological replicates, ribbons, SEM across replicates (WT, *n* = 11; F206D, *n* = 4; 4D, *n* = 3; L454K, *n* = 3 independent experiments). Lower strip: per-shell significance of each variant versus F206D; variants were compared across separate experiments by normalizing to each internal WT control (not shown). Tile color = −log₁₀(FDR), ** < 0.01, * < 0.05 by Welch’s unpaired *t*-test with BH correction. C) FACS plots of WT or lamin A F206D-expressing epicardial cells treated with vehicle or 5 µM MG132 for 18 hours, then stained for LMNA/C. Quantified in (Figure 3I).

**Figure 4 supplement 1.**
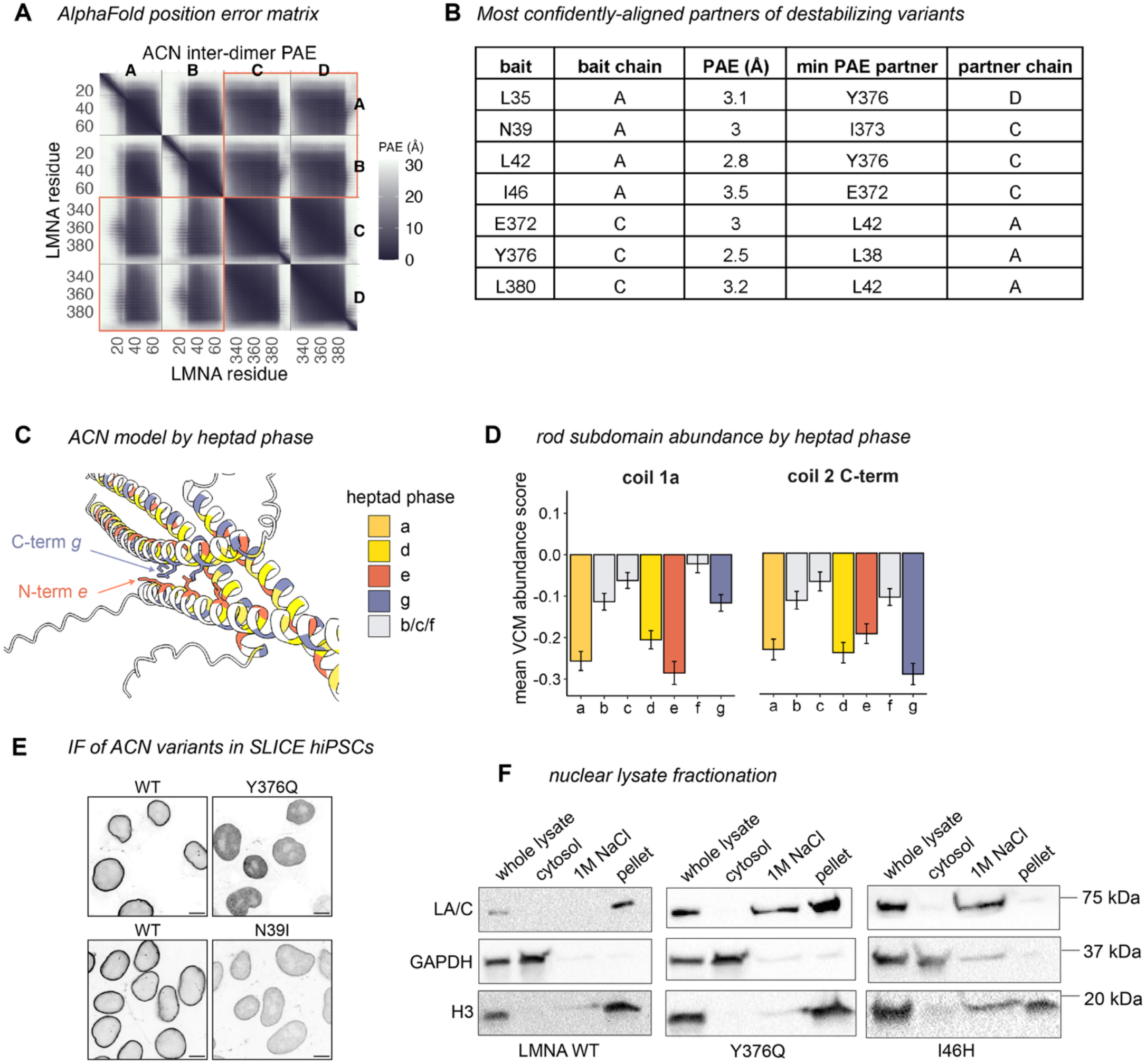
Terminal destabilizing residues constitute the ACN interface. A) Predicted alignment error (PAE) matrix for AlphaFold3 ACN model. N-terminal chains A/B and C-terminal chains C/D form contacts at upper right and bottom left (orange boxes). Residue positions for each chain indicated on the bottom and left axes. B) Table of predicted alignment error between destabilizing residues (bait) and their most confidently aligned partner. N-terminal dimer chains A/B and C-terminal dimer chains C/D for each residue indicated. C) AlphaFold3 ACN model colored by heptad phase. Orange N-terminal *e* phase face aligns with blue C-terminal *g* phase face, indicated by arrows. D) Bar plots of mean VCM protein abundance ± SD for each heptad phase in coil 1a (positions 28–70) and the C-terminus of coil 2 (positions 350–388). E) Immunofluorescence images of *LMNA* KO cells expressing indicated lamin A mutants in SLICE landing pad, stained with Ig-fold LMNA/C antibody. Scale bars, 10 µm. F) Western blots of high-salt nuclear fractionations. *LMNA* KO hiPSCs expressing the indicated variants from the SLICE landing pad were snap frozen and gently lysed by freeze/thaw. Nuclei were extracted with 1 M NaCl and separated into soluble and insoluble fractions, then blotted for LMNA/C, GAPDH (cytosolic marker) and histone H3 (nuclear marker).

**Figure 5 supplement 1.**
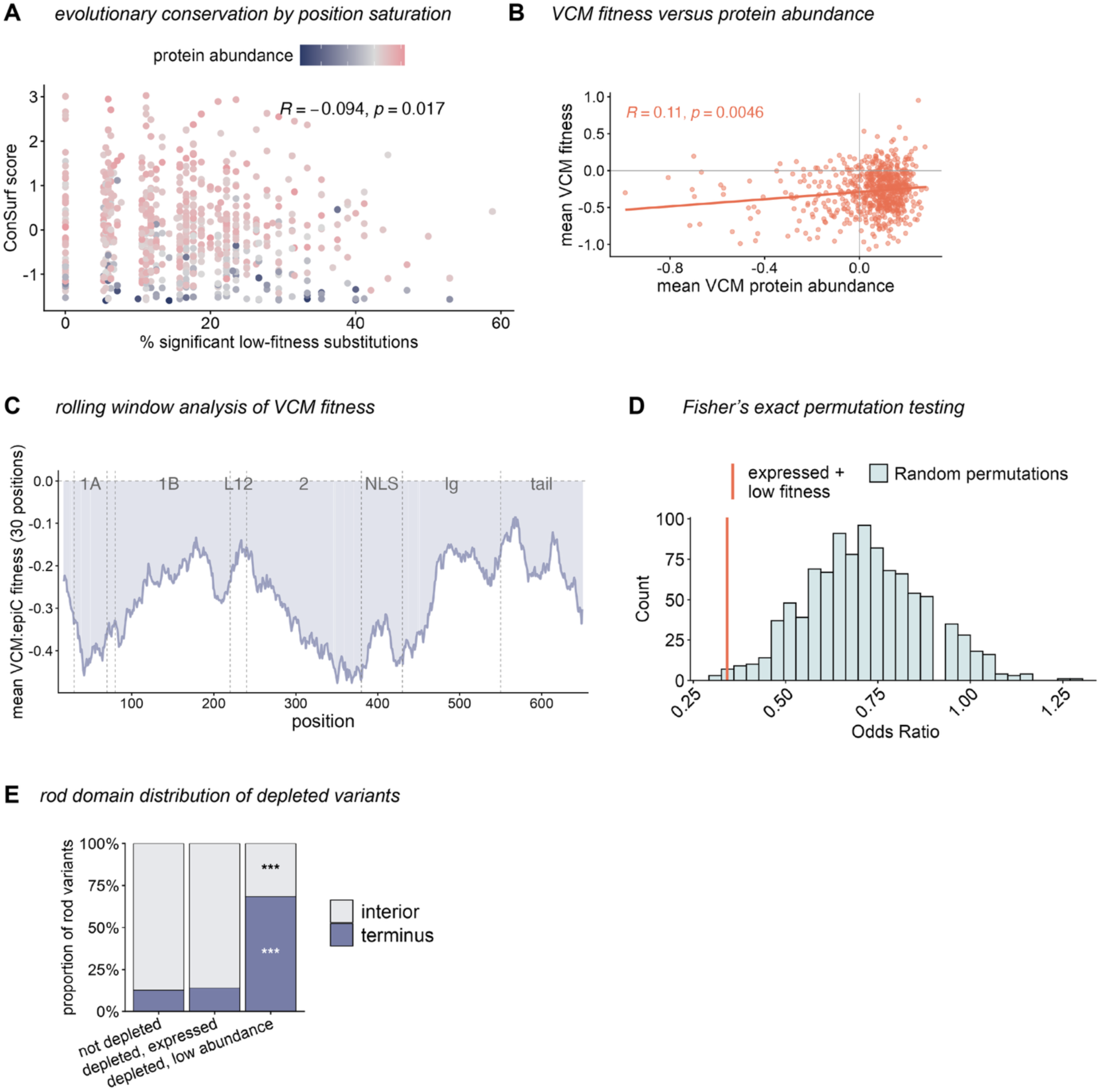
Conservation and location of low-fitness variants. A) Positions with more low-fitness variants are more evolutionarily conserved. x-axis: % of variants at each position called significantly depleted in VCMs relative to epiCs by Rosace. y-axis: ConSurf evolutionary conservation score, where more negative values are more conserved. Points are colored by VCM protein abundance. Top right, Spearman correlation. B) Correlation between VCM protein abundance and fitness scores. Upper left, Spearman correlation not subset by ClinVar status. C) Rolling means of 30-position windows across the lamin sequence, where each position represents the window center. Vertical dashed lines indicate domain boundaries and domains are labeled along the top. D) Companion figure to (Figure 5G). Odds ratio for presence in population databases (gnomAD + Regeneron) for the expressed low-fitness set (vertical orange line, see Methods) versus a null distribution of 1,000 mutation-type–matched random sets (histogram). Matching the random sets to the test set’s composition of missense/nonsense/synonymous variants controls for their differing population frequencies. The observed odds ratio (0.342) lies below the null (mean = 0.714, *P* = 0.011), indicating depletion from the population beyond expectation. E) Distribution of depleted variants at the rod domain interior versus terminus (first and last 25 amino acids). \*\*\**P* < 0.001: enrichment or depletion relative to the not-depleted set by Fisher’s exact test.

**Figure 6 supplement 1.**
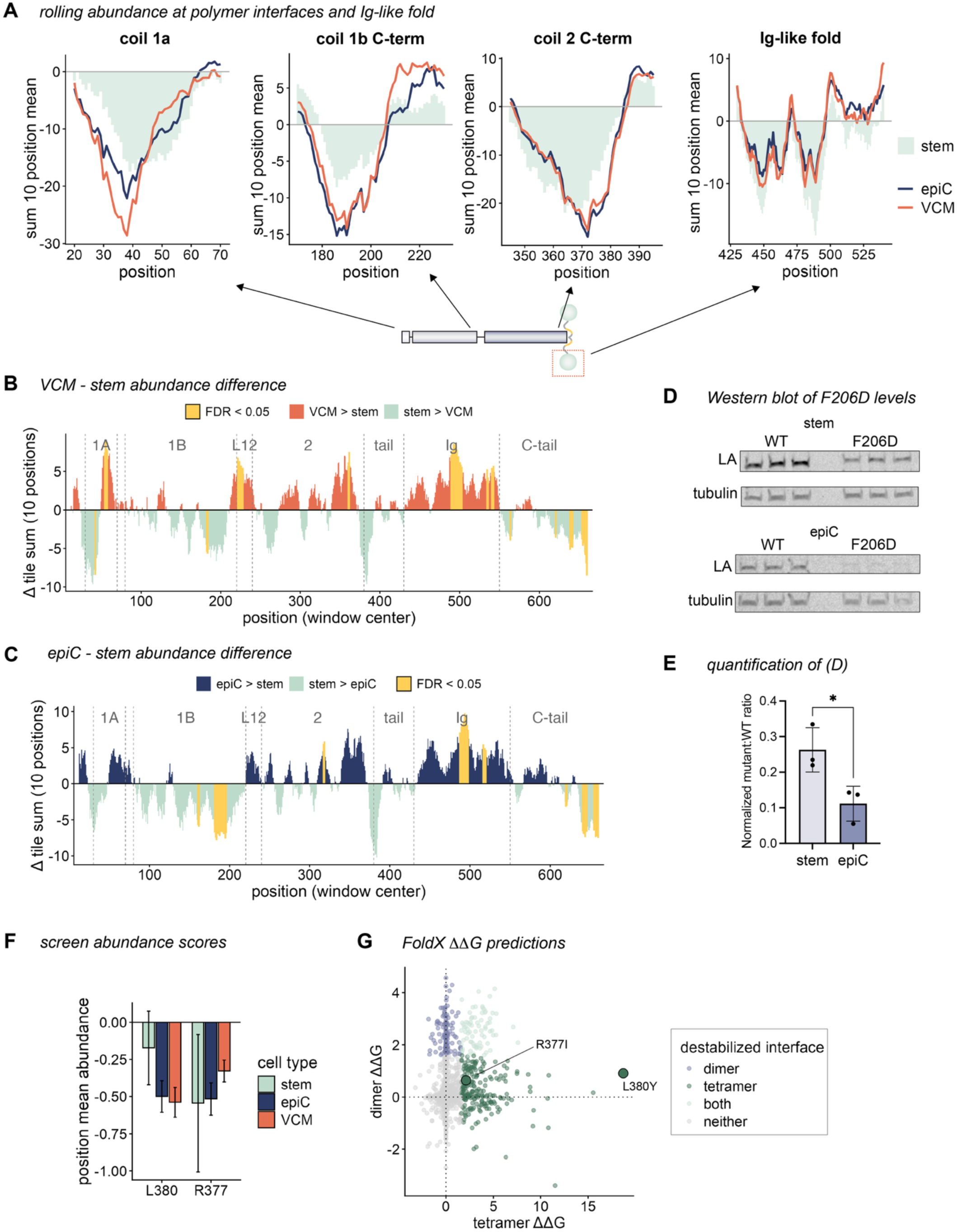
Polymer interface mutations are more destabilizing after cardiac differentiation. A) Sliding-window abundance signal within ACN, A11, and Ig-fold regions by cell type. Each trace is the rolling sum of z-scored position-mean Lilace abundance scores over 10-residue windows (1-position step), plotted at the window center, for each cell type (colors). Location of each region indicated on domain diagram below. B) Sliding-window comparison of position-level variant effects between VCMs and hiPSCs. Each bar is a 10-position sliding window (1-position step) along lamin A, plotted at the window center (x-axis). Bar height (Δ tile sum) = sum of the per-position z-score differences (Δ = z VCM − z hiPSC) across the 10 positions in each window. Positive values indicate higher lamin A abundance in VCMs, negative values indicate higher lamin A abundance in hiPSCs. Bar color denotes the direction of the difference; see key. Significance was assessed per window by a one-sample *t*-test of the 10 per-position differences against zero with BH FDR correction; windows with FDR < 0.05 highlighted in yellow. C) As in (B), sliding-window comparison of position-level variant effects between epiCs and hiPSCs. D) Western blot and E) quantification of *LMNA* KO stem cells and epicardial cells expressing lamin A F206D from SLICE landing pad. *n* = 3 independent experiments, triplicates run side-by-side. \**P* = 0.0323 by two-tailed Welch’s *t*-test. F) Positional mean abundance screen scores for SLICE-expressed L380 and R377 across cell types. Error bars = SD. G) FoldX mutational scan prediction for AlphaFold3 model ACN tetramer versus unit dimers. Points are colored by destabilized interface (variants with ΔΔG > 1.6 kcal/mol for tetramer, dimer, or both interfaces). R377I and L380Y, large points. Both are predicted to destabilize the tetramer, but L380Y has greater predicted energetic effect.

**Figure 6 supplement 2.**
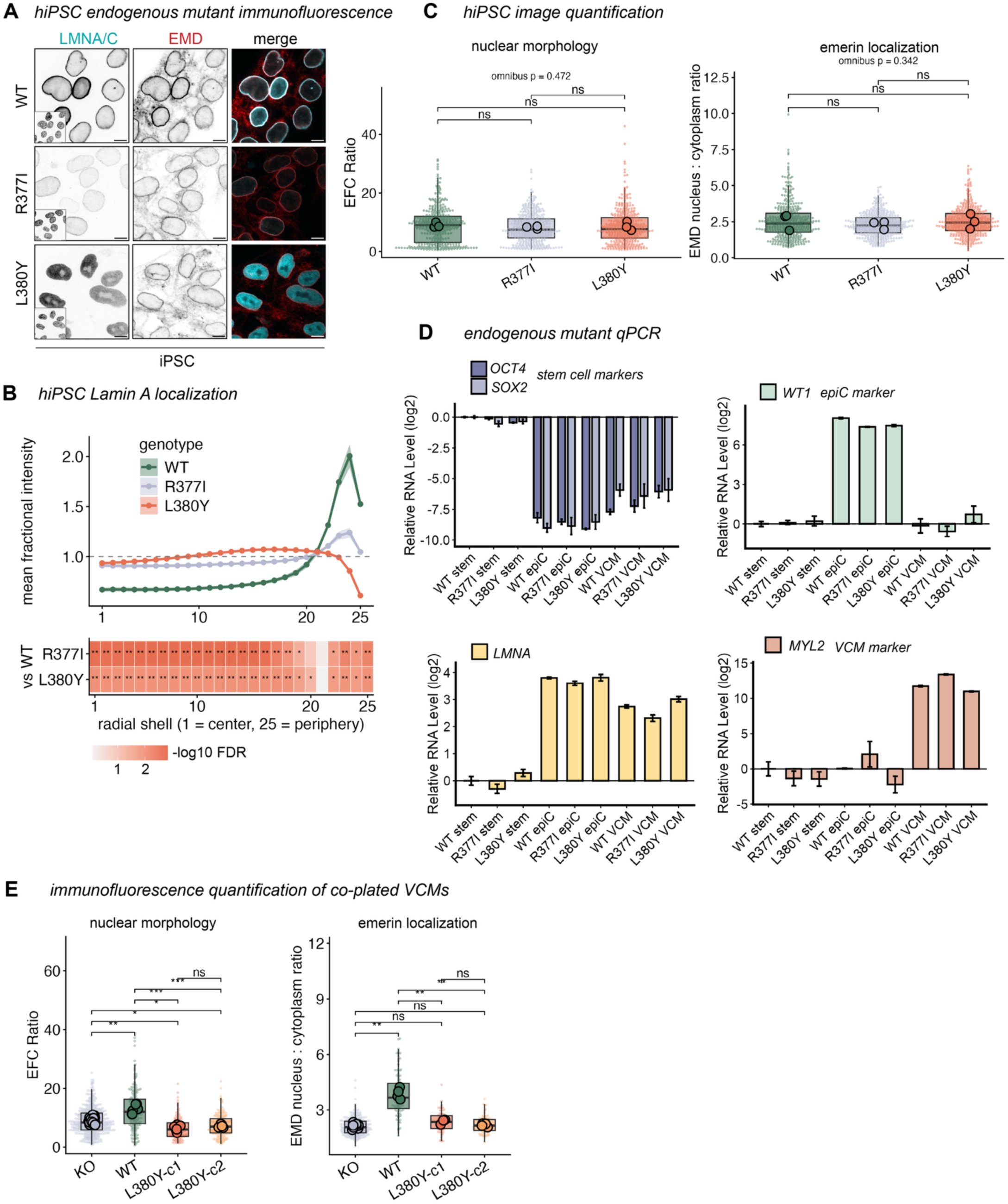
Distinct effects of R377I and L380Y variants across cell types. A) Immunofluorescence images of CRISPR-edited hiPSCs, fixed and stained for LMNA/C and emerin (EMD). Inset, Hoechst DNA stain. Scale bar, 10 µm. B) Quantification of hiPSC lamin A fractional intensity (shell mean ÷ whole-nucleus mean) from nuclear center (shell 1) to periphery (shell 25). Lines are means across biological replicates, ribbons, SEM across 3 independent experiments. Lower strip: per-shell significance of each variant versus WT. Tile color = −log₁₀(FDR), ** < 0.01, * < 0.05 by Welch’s unpaired *t*-test with BH correction. C) Box plots displaying hiPSC EFC ratio (left, higher = more regular nuclear morphology) and emerin nucleus:cytoplasm intensity ratio (right) across *LMNA* genotypes. Small points are individual nuclei (box = pooled median and IQR); large points, per-replicate means (the unit of *n*). Omnibus *P*: nested ANOVA (*n* = 3 independent experiments) indicates difference between all cell types. Brackets: Holm-corrected pairwise paired *t*-tests on replicate means (\**P* < 0.05, \*\**P* < 0.01, \*\*\**P* < 0.001). D) qPCR for *LMNA* transcript and lineage-specific genes in CRISPR-edited hiPSCs, epiCs, and VCMs bearing homozygous L380Y, R377I, or WT alleles. *OCT4* and *SOX2*: hiPSC markers, *WT1*: epicardial marker, *MYL2*: VCM marker. Transcript levels are normalized to WT hiPSC condition and log2 scaled. E) *LMNA* KO cells were co-plated with either WT or two separate L380Y clones, then fixed and stained for LMNA/C and emerin. Genotype assigned per nucleus from LMNA/C immunofluorescence, then EFC ratio (higher = more regular) and emerin nuclear enrichment was quantified using custom Python analysis pipeline. Small points are individual nuclei (KO *n* = 528, WT *n* = 206, L380Y-c1 *n* = 146, L380Y-c2 *n* = 204). Large outlined points are per-well means, on which all statistics were computed (*n* = 18, 6, 6, 6 wells), drawn from 4 independent platings across 2 differentiations and two differentiation timepoints (D29 and D32). KO shares a well with each expressing line, so KO-versus-line comparisons were made by paired *t*-test; WT and the L380Y clones were plated separately and never co-imaged, so comparisons among expressing lines were made by Welch’s unpaired *t*-test. \**P* < 0.05, \*\**P* < 0.01, \*\*\**P* < 0.001; ns, not significant.

